# Increased ambient temperature mitigates pathology in the *Ndufs4*^-/-^ mouse model of pediatric Leigh Syndrome

**DOI:** 10.1101/2025.10.24.684321

**Authors:** Melissa A.E. van de Wal, Christian J.M.I. Klein, Merel J.W. Adjobo-Hermans, Els M.A. van de Westerlo, Clara van Karnebeek, Mirian C.H. Janssen, Jan. C. van der Meijden, Judith R. Homberg, Jaap Keijer, Evert M. van Schothorst, Werner J.H. Koopman

**Affiliations:** Department of Pediatrics, Amalia Children’s Hospital, Radboud University Medical Center, Nijmegen, The Netherlands; Radboud Center for Mitochondrial Medicine, Radboud University Medical Center, Nijmegen, The Netherlands; Human and Animal Physiology, Wageningen University, Wageningen, The Netherlands; TSE Systems GmbH, Berlin, Germany; Department of Medical BioSciences, Radboud University Medical Center, Nijmegen, The Netherlands; Department of Pediatrics, Emma Personalized Medicine Center, Emma Children’s Hospital, Amsterdam University Medical Centers, Amsterdam, The Netherlands; Department of Internal Medicine, Radboud University Medical Center, Nijmegen, The Netherlands; Department of Cognitive Neuroscience, Donders Institute for Brain, Cognition and Behaviour, Radboudumc, Nijmegen, The Netherlands

**Keywords:** survival, thermoneutrality, thermoregulation, indirect calorimetry

## Abstract

Leigh syndrome (LS) is a devastating mitochondrial disease (MD) for which there is no treatment. Together with Leigh-like syndrome (LLS), LS constitutes part of the Leigh syndrome spectrum (LSS) disorders, which are the most frequent manifestation of a primary mitochondrial disease (MD) in children. *Ndufs4*^*-/-*^ mice are a widely used animal model to study LS pathophysiology and interventions. These mice display an isolated mitochondrial complex I deficiency and a brain-specific pathomechanism. Similar to other mouse models of human disease, *Ndufs4*^*-/-*^ mice are routinely housed and studied at a sub-thermoneutral ambient temperature. This means that these mice experience chronic cold-stress, which potentially aggravates disease symptoms and reduces their translational value. Here, we provide evidence that housing *Ndufs4*^*-/-*^ mice at 26 °C instead of 20 °C increases their skin, core and brain temperature. At this higher temperature, *Ndufs4*^*-/-*^ mice displayed lower energy expenditure and, importantly, a longer lifespan, pathology reversal in specific brain regions, as well as increased voluntary locomotor activity. We conclude that ambient temperature is a previously overlooked but highly relevant disease modifier in *Ndufs4*^*-/-*^ mice. Given the reduced mitochondrial energy production and aberrant thermoregulation in LSS and other MD patients, our findings suggest that reducing energy requirements might be of therapeutic value and/or contribute to an improved quality of life in these patients. In a broader sense, our results advocate the use of (more) thermoneutral housing to evaluate pathomechanisms and intervention strategies in murine models of human disease.

**Significance:** We conclude that ambient temperature is a previously overlooked but highly relevant disease modifier in *Ndufs4*^*-/-*^ mice. Given the reduced mitochondrial energy production and aberrant thermoregulation in LS and other MD patients, our findings suggest that reducing energy requirements might be of therapeutic value and/or contribute to an improved quality of life in these patients. In a broader sense, our results clearly demonstrate why it is essential to use a (more) thermoneutral housing to evaluate pathomechanisms and intervention strategies in translational research with murine models of human disease.

## INTRODUCTION

Leigh syndrome (LS; OMIM25600), or subacute necrotizing encephalomyelopathy, is a pediatric mitochondrial disease (MD) first described by Archibald Denis Leigh (**Leigh, 1951**). LS can be caused by gene mutations in the nuclear (nDNA) or mitochondrial DNA (mtDNA), which encode subunits or assembly factors of the oxidative phosphorylation (OXPHOS) system (**Koopman et al., 2013**). LS presents with a characteristic neuropathology, including multiple focal symmetric necrotic lesions in the brain stem, optic nerves, dentate nuclei, thalamus and basal ganglia (**Leigh, 1951; Baertling et al., 2014; Stenton et al., 2022; McCormick et al., 2023; Rahman, 2023; Thorburn et al., 2023; Ball et al., 2024**). At the histological level, these lesions appear spongiform and display vascular proliferation, gliosis and demyelination (**Rahman & Thorburn, 2020**). Analysis of 385 LS patients demonstrated that the most common clinical signs during disease progression are: developmental delay, hypotonia (low muscle tone and reduced muscle strength), respiratory dysfunction, epileptic seizures, poor feeding, and weakness (**Chang et al., 2020**). Together with Leigh-like syndrome (LLS), LS constitutes part of the Leigh syndrome spectrum (LSS) disorders, which are the most frequent manifestation of a primary MD in children (**McCormick et al., 2023**). LSS symptoms start occurring between 3-12 months of age with 50% of the patients dying by the age of 3 years, often due to cardiac or respiratory failure (**Rahman & Thorburn, 2020; Thorburn et al., 2023**). Due to an incomplete understanding of the LSS pathomechanism no treatments exist, and the prognosis is poor.

Deficiencies of mitochondrial complex I (CI), the first complex of the oxidative phosphorylation (OXPHOS) system (**Smeitink et al., 2001**), are the most frequent cause of LSS (**Rahman, 2024**). About 5% of the cases of nDNA-linked LSS are due to autosomal recessive mutations in the *NDUFS4* gene (**Rahman et al., 2023**). This gene encodes the NDUFS4 (NADH dehydrogenase [ubiquinone] iron-sulfur protein 4) protein, which is a CI structural subunit (**Papa et al., 1996; van den Heuvel et al., 1998; Budde et al., 2003; Anderson et al., 2008; Assouline et al., 2012; Assereto et al., 2014; Lamont et al., 2016; Ortigoza-Escobar et al., 2016**). Apparently, NDUFS4 plays a role in CI biogenesis and stabilization, thereby affecting CI levels and activity (**Kahlhöfer et al., 2017; Adjobo-Hermans et al., 2020; Yin et al., 2024**). At the protein level, pathogenic *NDUFS4* mutations induce complete absence of NDUFS4 protein, as well as of the CI structural subunit NDUFA12 (**Adjobo-Hermans et al., 2020**).

*Ndufs4* whole-body knockout (KO) mice (*Ndufs4*^*-/-*^ mice) represent a widely used animal model to study the LS pathomechanism and intervention strategies (**Kruse et al., 2008; Quintana et al., 2010; Quintana et al., 2012; Johnson et al., 2013; Jain et al., 2016; de Haas et al., 2017; Bolea et al., 2019; Martin-Perez et al., 2020; Henke et al., 2024; Blume et al., 2025**). Similar to LS patients with pathogenic *NDUFS4* mutations, *Ndufs4*^*-/-*^ mice exhibit brainstem and basal ganglia aberrations, ataxia and motor alterations, growth retardation, failure to thrive, hypotonia, visual problems, breathing irregularities and apneic episodes (**Quintana et al., 2012**). These mice develop a fatal encephalomyopathy and display a brain-specific phenotype leading to early death (**Kruse et al., 2008; Van de Wal, 2022**).

It is well recognized that the housing of mice at a sub-thermoneutral ambient temperature (T_A_) leads to cold-induced stress, which profoundly reduces their translational value in human disease modeling (**Tschöp et al., 2012; Abreu-Vieira et al., 2015; Hylander & Repasky, 2016; Ganeshan & Chawla, 2017; Gordon, 2017; Seelay & MacDougald, 2021; Zhao et al., 2022b; Hylander et et al., 2023; James et al., 2023; Lac et al., 2023; MacDonald et al., 2023**). For example, mice housed at 26 °C devote ∼20% of their total energy expenditure to cold-induced thermogenesis, but this number increases to ∼40% at 22 °C to maintain T_B_ (**Abreu-Vieira et al., 2015**). We previously argued that in order to better mimic human thermal relations and thereby improve the translational value of mouse models, these mice should be housed at 2-3 °C below their thermoneutral zone (*i*.*e*. below 28 °C; **Speakman & Keijer, 2012; Keijer & Speakman, 2019a; Keijer & Speakman, 2019b**). Evidence from the literature suggests that *Ndufs4*^*-/-*^ mice have a 2-4 °C lower T_B_ relative to control mice around postnatal day (PD) 33. Apparently, this temperature drop precedes development of severe symptoms and brain pathology, which occur between PD39-44 (**Fig. 1A**). Therefore, it is to be expected that a too low T_A_ is particularly relevant for *Ndufs4*^*-/-*^ mice and other MD mouse models with impaired mitochondrial ATP production. In healthy mice, T_A_ is a key determinant of energy balance, body mass and body composition (**Zhao et al., 2022a**), whereas T_B_ is an important modulator of lifespan (**Zhao et al., 2022b**). This suggests that increasing T_A_ and/or T_B_ might have beneficial effects in *Ndufs4*^*-/-*^ mice. Interestingly, integrating data from previous *Ndufs4*^*-/-*^ mouse studies revealed a positive exponential correlation between T_B_ and age of death (AOD; **Fig. 1B**). This suggests that T_B_ is a (co)modifier of the disease phenotype in these mice. In the current study we investigated this hypothesis and demonstrate that housing *Ndufs4*^*-/-*^mice at T_A_ = 26 °C instead of T_A_ = 20 °C increases their skin, core and brain temperature. These increases were paralleled by a lower energy expenditure, longer lifespan, reversal of pathology in specific brain regions, and increased voluntary locomotor activity. We conclude that T_A_ is an important modifier of the *Ndufs4*^*-/-*^ disease phenotype, potentially suggesting that increasing T_A_ and/or reducing heat loss might be of therapeutic value in LS/LSS and other MD patients.

**Figure 1:**
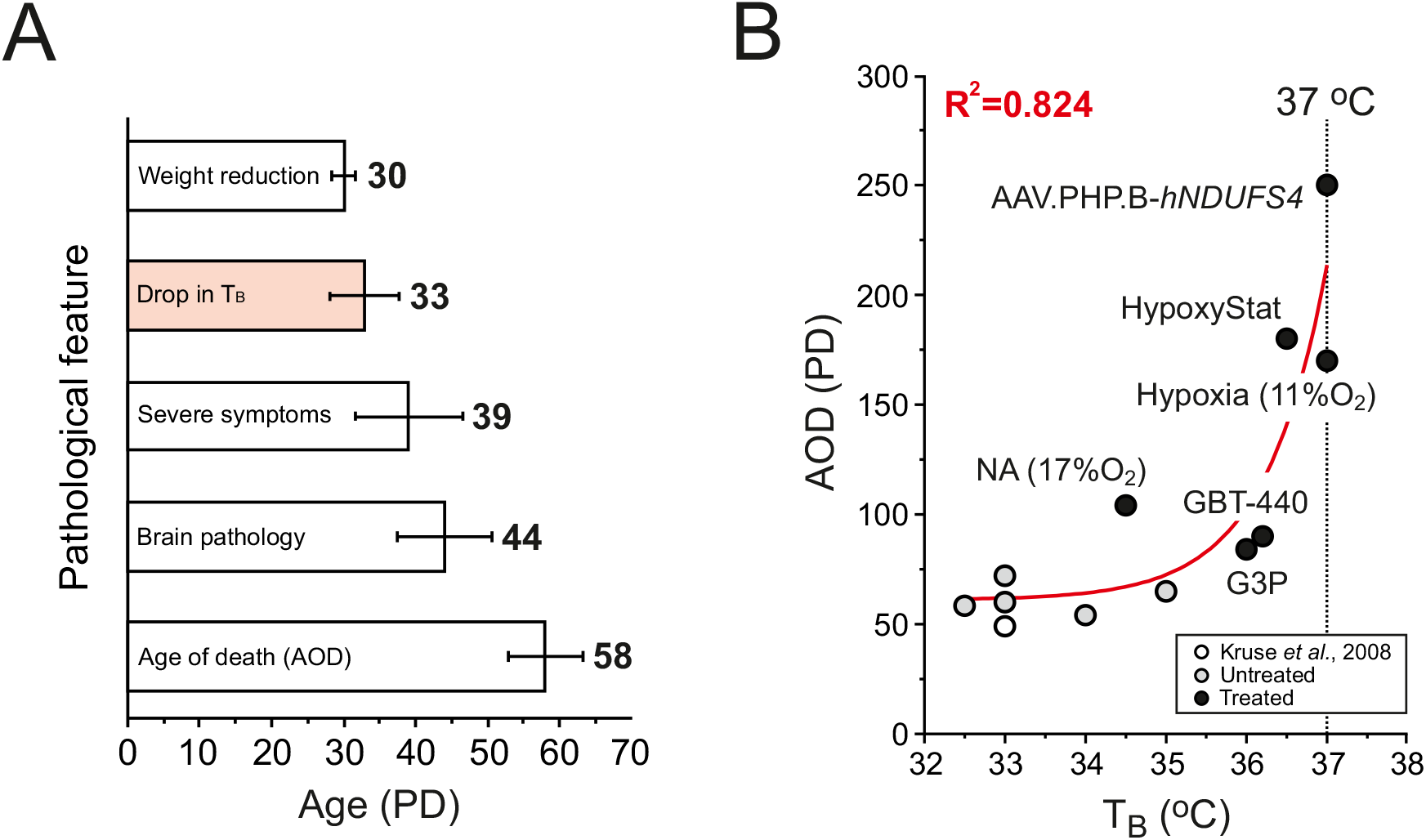
Disease symptom chronology, body temperature and age of death in KO mice. (**A**) Occurrence of key pathological features as a function of age (postnatal day; PD) in *Ndufs4*^*-/-*^ (KO) mice reported in the literature (**Supplementary Table S1**). T_B_ indicates body temperature. The number in bold indicates the mean age (PD) at which the feature occurs. (**B**) Correlation between T_B_ and age of death (AOD) of KO mice reported in the literature (**Supplementary Table S2**). The symbols reflect data obtained in untreated (gray) and intervention-treated (black) KO mice. The open symbol reflects data from the study first describing the KO model (**Kruse et al., 2008**). **Statistics:** In panel A, bars and errors reflect mean<u>±</u>standard deviation (SD). In panel B, a mono-exponential function was used for fitting (AOD=A·EXP(T_B_/µ)+y_0_), with A=2.4E-19±4.5E-18 (SE), µ=0.77±0.30 (SE) and y_0_=61±12 (SE). **Abbreviations:** AAV.PHP.B-*hNDUFS4* = adeno-associated virus (AAV)-delivered human *NDUFS4* (*hNDUFS4*); G3P = glycerol-3-phosphate; GBT0440 (Voxelotor) = a compound for sickle cell anemia; NA = nicotinic acid.

## METHODS

### Animals

Heterozygous mouse breeding pairs (*Ndufs4*^*+/-*^; B6.129S4-Ndufs4tm1.1Rpa/J; #027058) were purchased (**Jackson Laboratory; Bar Harbor, Maine, USA**) and intercrossed to generate *Ndufs4*^*-/-*^ whole-body knockout (KO) and wildtype (WT) littermate mice as described previously (**Kruse et al., 2008; de Haas et al., 2016**). Voluntary locomotor analysis and longevity studies were performed at the Animal Research Facility of the Radboud University Medical Center (**Radboudumc, Nijmegen, The Netherlands**) as follows: mice were group-housed under controlled conditions including bedding, nesting material, shelter and cage enrichment at an ambient temperature (T_A_) between 20-21°C in Makrolon (polycarbonate) Type II cages (humidity 50-70%; 12 h light/dark cycle; lights on at 8 AM). At postnatal day (PD) 8-9, toe clips were taken for identification and genotyping. At PD21, experimental mice (n=15 for each group) were weaned and allocated to be group-housed per sex at T_A_ = 20 °C or T_A_ = 26 °C. Mice had *ad libitum* access to food and water and were fed a standard animal diet (V1534-300 R/M-H; **Ssniff GmbH, Soest, Germany**). KO mice were maintained until the humane endpoint was reached. Whole-body energy metabolism and -related analyses (see below) were performed at the Carus Animal Research Facility of Wageningen University (**WU, Wageningen, The Netherlands**). Mice were housed at T_A_ = 23 °C under controlled conditions, including bedding, nest material and cage enrichment (humidity 50±10%; 12 h light/dark cycle; lights on at 5 AM) in Makrolon (polycarbonate) Type II cages and, unless stated otherwise, had *ad libitum* access to a standard chow diet (Teklad Global Diet 2920; **Inotiv Inc., IN, USA**) and water. Pups that were born on the same day (±1 day), were considered to be of the same age and used as experimental mice. At PD9, toe clips were taken for identification and genotyping. Nests remained with their dam (female breeder) until PD21. At PD21, mice were weaned and experimental mice (n=8 per group) were individually housed at either T_A_ = 20 °C or T_A_ = 26 °C. Animal studies were conducted under an ethical license provided by the national competent authority (CCD, Centrale Commissie Dierproeven, AVD10300-20173-827 at **Radboudumc**, AVD10400-20231-7449 at **WU**), following a positive advice from its local Animal Ethics Committee (DEC). All animal procedures were captured in protocol and approved by the Animal Welfare Bodies (2017-0017-007 at **Radboudumc** and 2023.W-0017.001 at **WU**), securing full compliance to the European Directive 2010/63/EU for the use of mice for scientific purposes.

### Voluntary locomotor analysis

Individual male/female KO and WT mice were used for voluntary locomotor analysis (total duration: 23 h; light phase: 11 h; dark phase: 12 h) on PD25, PD30, PD35 and PD40. This was performed using a PhenoTyper system (**Noldus Information Technology, Wageningen, The Netherlands**), as described (**de Visser et al., 2006**). Briefly, each cage contained a top unit with a digital infra-red (IR) video camera, equipped with an IR filter to prevent interference of ambient light during the light phase, and IR lighting. Cages consisted of transparent polymethyl methacrylate (Perspex) walls with an aluminum floor and had dimensions of 45(L) x 45(W) x 55(H) cm. The floor was covered with bedding without nesting material to prevent interference with positional animal tracking. Food and water were available *ad libitum* at the bottom of the cage. A shelter was placed in one of the corners of the cage. For each animal, the total distance traveled was quantified using Ethovision XT software (version 15.1; **Noldus Information Technology**).

### Collection of brain tissue

Once the humane endpoint of the KO mice used for voluntary locomotor analysis was reached, WT and KO mice were sacrificed by decapitation. After dissection, a batch of whole brains (n=7-8) was stored (−80 °C) for Western blot analyses. For immunocytochemistry, a second batch (n=7-8) was fixed in ice-cold 4% (v/v) paraformaldehyde (pre-dissolved in PBS overnight) containing 30% (w/v) sucrose (#57-50-1; **Thermo Fisher Scientific, Waltham, MA, United States**). These brains were cryopreserved by storing them in PBS containing 30% (w/v) sucrose (**Thermo Fisher Scientific**) for 2-5 days.

### Western blotting

Half brains (right hemisphere) were incubated (30 min, on ice) in RIPA buffer (50 mM NaCl, 1% (w/v) Triton X-100, 5 mM Na_2_EDTA, 10 mM Na_4_P_2_O_7_· 10H_2_O, 50 mM NaF, 50 mM Tris-HCl pH 7.4), containing 0.1 mg/ml DNAse (#79254; **Qiagen, Hilden, Germany**). Next, samples were centrifuged (11,000 x *g*; 4 °C; 10 min). Protein concentrations were determined using a Protein Assay Dye Reagent Concentrate (#500-0006; **Bio-Rad Laboratories B.V., Lunteren, The Netherlands**), according to the manufacturer’s instructions. Spectrophotometric absorbance was measured at 595 nm using a Benchmark Plus plate reader (**Bio-Rad**). Proteins (50 µg/lane) were separated on a 4-15% gradient SDS-PAGE gel (#4561086, **Bio-Rad**). Molecular weight (MW) markers were included for each gel (Precision Plus Protein Dual Color Standard; #1610374; **Bio-Rad**).Then, proteins were transferred to a methanol pre-wet PVDF membrane (#IPVH00010; **Merck-Millipore, Burlington, MA, USA**), incubated (1 h; RT) with blocking buffer (#927-70001, **Li-COR Biosciences, Lincoln, NE, USA**), and stained with primary antibodies (overnight at 4 °C). Antibodies included: cold-inducible RNA-binding protein (CIRBP; 1:1,000; #10209-2-AP; **Proteintech Europe, Manchester, UK**), RNA-binding motif protein 3 (RBM3; 1:1,000; #14363-1-AP; **Proteintech**), the complex I subunit NDUFB8 (1:1,000; #ab110242; **Abcam, Cambridge, UK)**, the complex II subunit SDHA (1:1,000; #ab14715; **Abcam**), the complex III subunit UQCRC2 (1:1,000 #ab14745; **Abcam**), the complex IV subunit MtCOX2 (1:2,000; #55070-1-AP; **Proteintech**), the complex V subunit ATP5A (1:1,000; #ab14748, **Abcam**) and β-actin (loading control; 1:100,000; #A5441; **Sigma-Aldrich, St. Louis, MO, USA**). Next, membranes were washed with 0.1% (v/v) Tween-20 in PBS (PBST) and incubated (1 h; RT) with the secondary antibodies GaRabbit IRdye800 (1:10,000; #92632211; **LI-COR Biosciences, Lincoln, NE, USA**) and GaMouse (1:10,000; IRdye680; #926-32220; **LI-COR**). Finally, blots were washed in PBTS, subsequently scanned on an Odyssey CLx imaging system (**LI-COR**), and images were stored in TIFF format. Quantification was performed using FIJI freeware (version: 1.54k). Protein levels were normalized on β-actin and expressed as percentage of the average value obtained with WT samples on the same blot.

### Immunocytochemistry

Brain sections (30 µm thick) were obtained from the cryopreserved brains using a microtome (Microm HM 440 E; **GMI, Bunker Lake, MN, USA**). These sections contained various brain regions (*e*.*g*. hippocampus, olfactory bulb, motor cortex and superior colliculus) and were stored in 0.01% (v/v) sodium azide in PBS at 4°C. Sections were stained using a free-floating method (**Kelly et al., 2022**). Briefly, slices were washed in PBS (3 times for 5 min) and then placed in a blocking buffer (PBS containing 10% (w/v) BSA and 0.3% (v/v) Triton-X-100) for 30 min at RT. Next, slices were incubated overnight at 4 °C with antibodies against the microglia marker ionized calcium-binding adapter molecule 1 (IBA1; a.k.a. Allograft Inflammatory Factor 1; AIF1; 1:1000, #234017; **Synaptic Systems, Göttingen, Germany**) and the astrocyte marker glial fibrillary acidic protein (GFAP; 1:1000, #173006; **Synaptic Systems**). Then, the slices were washed in PBS (three times for 10 min) followed by incubation with fluorescent secondary antibodies (3 h; RT) for detection of IBA1 (goat anti-chicken Alexa Fluor 594; 1:2,000, #A11042; **Thermofisher Scientific**) and GFAP (goat anti-rat Alexa Fluor 488; 1:2,000, A21212; **Thermofisher Scientific**). DNA staining was carried out during the last 15 min of the secondary antibody incubation by adding 4′,6-diamidino-2-phenylindole (DAPI; 1.67 µg/ml; #32670; **Sigma-Aldrich**). Next, the slices were washed in PBS (three times for 5 min) and mounted on gelatin-coated microscopy slides (#S8902; **Sigma Aldrich**). Fluorescence microscopy was performed using an Axio Observer 7 system equipped with an x10 objective and AI sample finder (**Carl Zeiss AG, Oberkochen, Germany**). Image stacks (16 bit; three fluorescence channels: DAPI, IBA1, GFAP) were stored in CZI format and converted to TIFF stacks using FIJI freeware (version: 1.54k). IBA1- and GFAP-positive cells were manually counted in regions of interest (ROIs), defined using the Allen brain atlas (https://mouse.brain-map.org). For normalization, the number of positive cells was expressed per cm^2^.

### Whole-body energy metabolism and -related analyses

From PD14 until PD36 mice were weighed daily at the same time (morning). From PD26 to PD36, skin temperature (T_S_) was monitored daily using a RS PRO 8861 infrared (IR) thermometer (#303-91-528; **RS components, Haarlem, The Netherlands**). This thermometer was pointed at the perianal region until both beams converged in a single point to measure T_S_. Animal core temperature (T_C_) was calculated using an empirical equation (T_C_=-0.12·(T_S_-30)^2^+0.8·(T_S_-30)+26.36; **Mei et al., 2018**). To determine lean mass (LM) and fat mass (FM), a body composition analysis (BCA) was performed at PD22, PD29, PD30, PD34 and PD36 using an Echo-MRI Whole Body Composition Analyzer (**Echo-MRI, Houston, FL, USA**). Between PN30 and PN34, male mice were individually placed in an indirect calorimetry (InCa) system (see below), continuing their T_A_ = 20 °C or T_A_ = 26 °C. Each cage (Makrolon IVC Green line; **Tecniplast S.p.A., Buguggiate, Italy**) had dimensions of 37.1(L) x 18.9(W) x 12.6(H) cm, and contained bedding, reduced nest material (to prevent interference with activity measurements), and cage enrichment. Day 1 and 2 (PD30-PD31) of the InCa experiments were used for acclimatization. InCa measurements were performed on day 3 (PD32) using a PhenoMaster Phenotyping Research Platform (**TSE Systems GmbH, Berlin, Germany**) as published previously (**Fernández-Calleja et al., 2018; Fernández-Calleja et al., 2019**), with the following changes: (**a**) the system contained 24 instead of 12 cages, (**b**) all gas sensor units were duplicated to keep the same sample interval. The InCa system employed dual units for gas analysis (**ABB Automation, Frankfurt am Main, Germany**) using internal calibration cells for ^12^CO_2_ and ^13^CO_2_ with dual air-drying units. A single reference cage was used across both dual systems. Measurements were recorded at intervals of 1.32 minutes per cage, resulting in three data points per h for each cage. Before each measurement run, gas sensor units were calibrated using internal calibration cells and specific calibration gasses (**Linde Gas Benelux BV, Dieren, The Netherlands**): zero (20.947% O_2_ together with solely N_2_), and span (0.5% O_2_, together with solely N_2_) gasses. The respiratory exchange ratio (RER) was obtained by dividing the net CO_2_ production rate (V_CO2_, equaling the sum of ^12^CO_2_ and ^13^CO_2_ production) by the net oxygen consumption rate (V_O2_). The raw energy expenditure (“raw EE”; kcal/h) was computed using the Weir equation. For animal welfare reasons, KO mice received additional food pellets and a small Petri dish containing water on the bottom of the cage. This precluded automated quantification of food and water intake during InCa measurements. Therefore, the weekly food intake (FI) for each animal was determined by weighing between PD22-29 and PD29-36.

### Data analysis

Statistical analysis and curve fitting were performed using OriginPro (version 2025a; **OriginLab Corporation, Northampton, MA, USA**) and GraphPad Prism software (version 9.5.1; **GraphPad Software Inc., Boston, MA, USA**). The coefficient of determination (R^2^) was used to determine the quality of exponential fitting, with an R^2^ value of 1 indicating a perfect fit (**de Groof et al., 2000**). Survival curves were compared using a Gehan-Brelow-Wilcoxon test. Unless stated otherwise, data from multiple experiments is presented as mean±standard error of the mean (SEM) and statistical significance was evaluated using a non-parametric Mann-Whitney U test.

## RESULTS

### Effect of increased ambient temperature on lifespan, body weight, and brain levels of cold-induced proteins and OXPHOS subunits

This study investigates the hypothesis that increased ambient temperature (T_A_) is a mitigating disease modifier in a mouse model of human Leigh syndrome (LS). To this end (**Fig. 2A**), we analyzed wildtype (WT) and whole-body *Ndufs4* knockout (KO) mice housed at T_A_ = 20 °C until PD21, after which they were divided into two groups and housed at T_A_ = 20 °C (WT20 and KO20 condition) or T_A_ = 26 °C (WT26 and KO26 condition). The lifespan of KO26 mice was significantly longer than of KO20 mice (age of death (AOD) = 60.9±1.59 days *vs*. 53.4±1.46 days, p<0.01), with median AOD values of 51 days (KO20) and 61 days (KO26; **Fig. 2B**). AOD values of individual mice ranged between 47-66 days (KO20) and 51-72 days (KO26), demonstrating a maximum AOD increase of 53% in KO26 mice. No significant differences in AOD were observed between male (M) and female (F) mice (**Fig. 2C**). Bodyweight (BW) was lower in KO relative to WT mice (**Fig. 2D**), confirming previous findings (reviewed in: **van de Wal et al., 2022**). Maximum BW was similar in KO20 and KO26 mice, meaning that the higher T_A_ did not affect this parameter (**Fig. 2D**). Female WT20 and WT26 mice reached a lower maximum BW than the corresponding male mice (**Fig. 2D**). In contrast, KO20 and KO26 mice did not display such a sex difference (**Fig. 2D**). Compatible with their increased AOD (**Fig. 2B**), KO26 mice reached their maximum BW at a later age than KO20 mice (**Fig. 2E**). This age was similar for male and female mice (*i*.*e*. between PD40-42). It was previously demonstrated that hypothermia induces upregulation of cold-inducible RNA-binding protein (CIRBP) and RNA-binding motif 3 (RBM3; **Rosenthal et al., 2019; Tong et al., 2013**). In case of RBM3, its cold-dependent upregulation was already induced by a drop in temperature from 37 °C to 36 °C in neuronal and astrocyte cultures, demonstrating the exquisite temperature sensitivity of this mechanism (**Jackson et al., 2015**). Proteome analysis demonstrated that *Ndufs4*^*-/-*^ mice at PD42 displayed higher CIRBP and RBM3 levels in brain, liver, heart, kidney, diaphragm and skeletal muscle relative to WT mice (**Supplementary Fig. S1** and **Supplementary Table S3**). This supports the conclusion that *Ndufs4*^*-/-*^ mice display a lower systemic temperature. Next, we analyzed the effect of increased T_A_ on brain CIRBP and RBM3 protein levels as a proxy of brain temperature. Confirming our previous brain proteome analysis (**van de Wal et al., 2025**), CIRBP and RBM3 levels were increased in KO20 relative to WT20 brains (**Fig. 2F-G**). WT26 mice displayed lower CIRBP levels than WT20 mice, and expression of CIRBP and RBM3 was apparently reduced in KO26 relative to KO20 mice (**Fig. 2F-G**). This suggests that WT26 and KO26 mice have a higher brain temperature than WT20 and KO20 mice. Given the temperature-sensitivity of the OXPHOS system (**Ali et al., 2010; Pamenter et al., 2018; Von Schulze et al., 2021; Davis et al., 2023; Jørgensen et al., 2023; Shin et al., 2024**), we next determined whether T_A_ elevation affected the brain expression of OXPHOS subunits. Compatible with our previous proteome studies (**Adjobo-Hermans et al., 2020; van de Wal et al., 2025**), KO20 mice displayed lower levels of the CI subunit NDUFB8 relative to WT20 mice (**Fig. 2H-I**). In contrast, expression of subunits of CII (SDHA), CIII (UQCRC2), CIV (MtCOX2) and CV (ATP5A) were not affected in KO20 mice (**Fig. 2H-I**). NDUFB8 expression was not altered in KO26 brain, whereas SDHA levels were slightly increased in WT26 relative to WT20 mice (**Fig. 2H-I**). Collectively, these results demonstrate that: (**1**) KO26 mice live longer than KO20 mice, (**2**) KO26 mice likely have a higher brain temperature than KO20 mice, and (**3**) KO26 and KO20 mice display a similar OXPHOS subunit expression profile.

**Figure 2:**
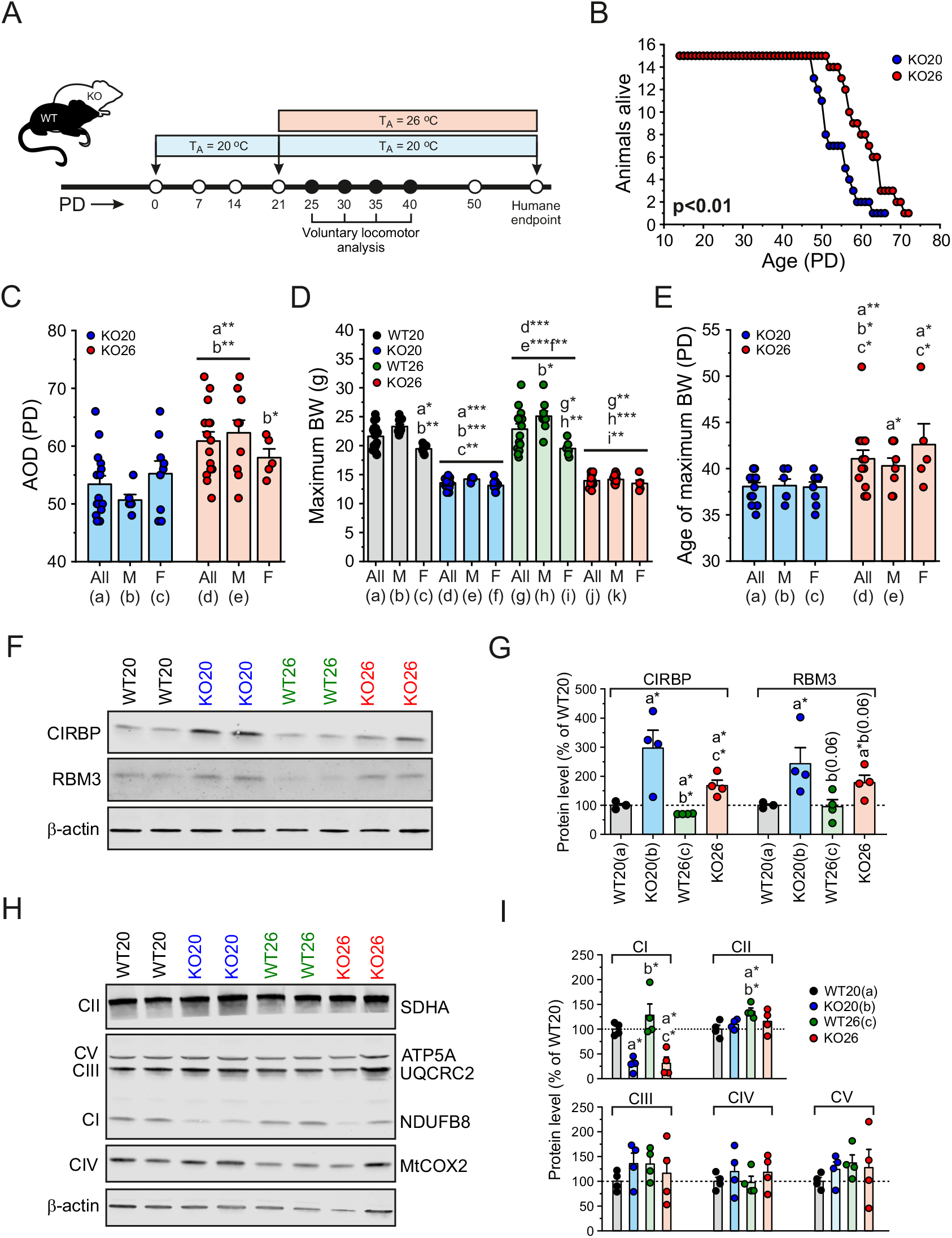
Effect of increased ambient temperature on the age of death of KO mice and the brain levels of cold-induced proteins and OXPHOS subunits in WT and KO mice. (**A**) Design of the thermo-intervention study and voluntary locomotor analysis. (**B**) Survival of KO mice housed at an ambient temperature (T_A_) of 20 °C (KO20) or 26 °C (KO26). (**C**) Age of death (AOD) expressed as postnatal days (PD) of KO20 and KO26 mice. Data is presented separately for all mice (“All”; gray bars), male mice (M), and female mice (F). (**D**) Maximum body weight (BW) of WT and KO mice housed at T_A_=20 °C (WT20, KO20) or 26 °C (WT26, KO26). Data is presented separately for all mice (all; gray bars), male mice (M), and female mice (F). (**E**) Age at which the maximum BW was reached for KO20 and KO26 mice. Data is presented separately for all mice (all; gray bars), male mice (M), and female mice (F). (**F**) Typical Western blot depicting expression of the cold-induced proteins CIRBP and RBM3 in brain homogenates of WB20, KO20, WT26 and KO26 mice. (**G**) Quantification of CIRBP and RBM3 Western blot data. (**H**) Same as panel F, but now for subunits of OXPHOS complex I to V (CI-CV). (**I**) Quantification of OXPHOS complex Western blot data. **Statistics:** Survival curves (panel B) were compared using a Gehan-Breslow-Wilcoxon test. Data in panels C, D and E was obtained for n=16 mice (WT20; 9 male, 7 female), n=15 (KO20; 6 male, 9 female), n=15 (WT26; 9 male, 6 female) and n=15 (KO26; 10 male, 5 female). Western blotting was performed using samples from n=3-4 mice (symbols) for each condition (N=2 independent blots). Western blot data was normalized on β-actin and expressed as percentage of the average WT20 value measured on the same blot. Western blot images were contrast optimized for visualization purposes (quantification was performed on the original blots; see **Supplementary Figure S2**). All significant differences between the indicated conditions (a,b,c,d,e,f,g,h,i,j,k) are marked by: *p<0.05, **p<0.01, ***p<0.001 or the exact p-value (borderline significance).

### Effect of increased ambient temperature on brain inflammation markers

Inflammation of lesion-prone brain regions, including the hippocampus and olfactory bulb, have previously been proposed as major contributors to the clinical phenotype and death of *Ndufs4*^*-/-*^ mice (**Johnson et al., 2012; Shil et al., 2021; Terburgh et al., 2021; Aguilar et al., 2022; Hanaford & Johnson, 2022; Stokes et al., 2022; van de Wal et al., 2022**). Moreover, mitochondrial dysfunction in the hippocampus is linked to psychiatric symptoms in MD patients (**Anglin et al., 2012**), and involvement of the motor cortex is suggested by the fact that KO mice display aberrant motor function and gait alterations (**Johnson et al., 2012; Quintana et al., 2012; de Haas et al., 2016; Schirris et al., 2021; Corrà et al., 2022**). We recently analyzed the proteomes of WT20 and KO20 mice with respect to the cerebellum, cerebral cortex, hippocampus, inferior colliculus and superior colliculus. This demonstrated that the largest number of proteome changes in KO20 mice occurred in the superior colliculus (**van de Wal et al., 2025**). In light of the above, we here determined whether an increased T_A_ altered neuroinflammation markers in the hippocampus, olfactory bulb, motor cortex and superior colliculus of WT and KO mice (**Fig. 3**). The inflammation marker proteins analyzed were: ionized calcium-binding adapter molecule 1 (IBA1; a.k.a. Allograft Inflammatory Factor 1 or AIF1; a microglia marker) and glial fibrillary acidic protein (GFAP; an astrocyte marker). These markers are commonly used to demonstrate microglia/astroglia activation and neuroinflammation (**Johnson et al., 2012; Quintana et al., 2012; Kempuraj et al., 2024; Naveed et al., 2024**). In the hippocampus, the number of IBA1-positive cells was similar between WT20, KO20 and WT26 mice, but reduced in KO26 mice (**Fig. 3A-B**). With respect to GFAP, hippocampal signals were increased in KO20 mice and apparently normalized in KO26 mice (**Fig. 3A-B**). In the olfactory bulb, relative to WT20 mice, the number of IBA1-positive cells were increased in KO20, WT26 and KO26 mice (**Fig. 3C-D**). GFAP signals were higher in the olfactory bulb of KO20 and KO26 mice, when compared to WT20 and WT26 mice (**Fig. 3C-D**). For the motor cortex, IBA1 signals were similar for WT20, KO20, WT26 and KO26 mice (**Fig. 3E-F**). In contrast, the number of GFAP-positive cells in the motor cortex was increased in KO20 mice and apparently normalized in KO26 mice (**Fig. 3E-F**). In case of the superior colliculus, no differences in IBA1 and GFAP signals between the four conditions were detected (**Fig. 3G-H**). Overall, these data demonstrate that: (**1**) KO20 mice display increased GFAP signals in the hippocampus, olfactory bulb and motor cortex, and (**2**) these GFAP signals are reduced in the hippocampus and motor cortex of KO26 mice.

**Figure 3:**
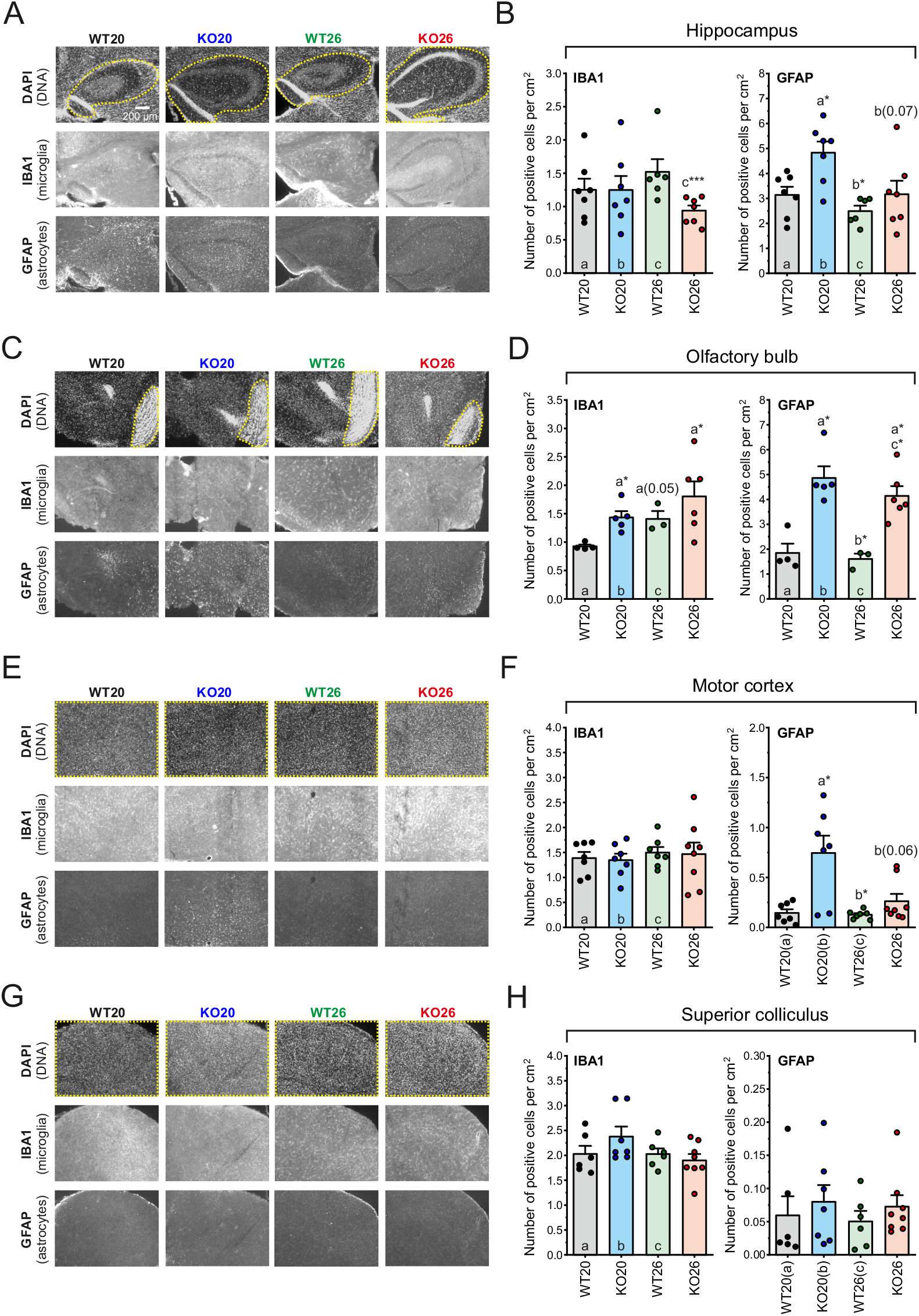
Effect of increased ambient temperature on pathology markers in specific brain regions of WT and KO mice. (**A**) Representative examples of hippocampal brain slices stained with DAPI (DNA) and antibodies against IBA1 (activated microglia) and GFAP (astrocytes). Samples were obtained from WT and KO mice housed at an ambient temperature (T_A_) of 20 °C (WT20, KO20) or 26 °C (WT26, KO26). The region of interest (ROI) used for quantification is highlighted in the DAPI image (yellow). (**B**) Average number of IBA1 and GFAP positive cells per cm^2^ in the hippocampal slice. (**C**) Same as panel A, but now for the olfactory bulb region. (**D**) Same as panel B, but now for the olfactory bulb region. (**E**) Same as panel A, but now for the motor cortex region. (**F**) Same as panel B, but now for the motor cortex region. (**G**) Same as panel A, but now for the superior colliculus region. (**H**) Same as panel B, but now for the superior colliculus region. **Statistics:** Microscopy images were contrast optimized for visualization purposes (quantification was performed on the original images). Data in panels B, D, F and H was obtained from multiple mice (symbols; n=3-8 mice for each condition). All significant differences between the indicated conditions (a,b,c) are marked by: *p<0.05 or the exact p-value (borderline significance).

### Effect of increased ambient temperature on voluntary movement activity

Given the involvement of the motor cortex in motor control (**Lee et al., 2022**), combined with the normalization of increased GFAP signals in this brain region (KO20) at increased T_A_ (KO26; **Fig. 3F**), we next quantified voluntary movement activity using automated animal tracking. To retain sufficient mice for analysis, their activity was studied during disease progression (PD25, PD30, PD35 and PD40), but prior to KO death (**Fig. 2B**). Because the effects of mitochondrial dysfunction in KO mice might have a greater impact in active mice, in addition to the total time period (23 h), data was also analyzed separately for the light (less active) and dark (more active) phase. In general, KO mice displayed lower total (**Fig. 4A**), light phase (**Fig. 4B**) and dark phase (**Fig. 4C**) activity relative to WT mice. Both WT and KO mice were more active at increased T_A_ (**Fig. 4A-B-C**). Activity increased for WT20 and WT26 mice as a function of age (PD), but not for KO20 and KO26 mice (**Fig. 4D**). Separate analysis of the light and dark phase revealed that WT20 and WT26 mice became more active in dark, whereas KO20 and KO26 mice did not (**Fig. 4E**). Taken together, these data demonstrate that: (**1**) the voluntary movement activity of WT and KO mice is increased at higher T_A_, (**2**) this activity increases a function of age for WT but not KO mice, and (**3**) WT but not KO mice become more active during the dark phase.

**Figure 4:**
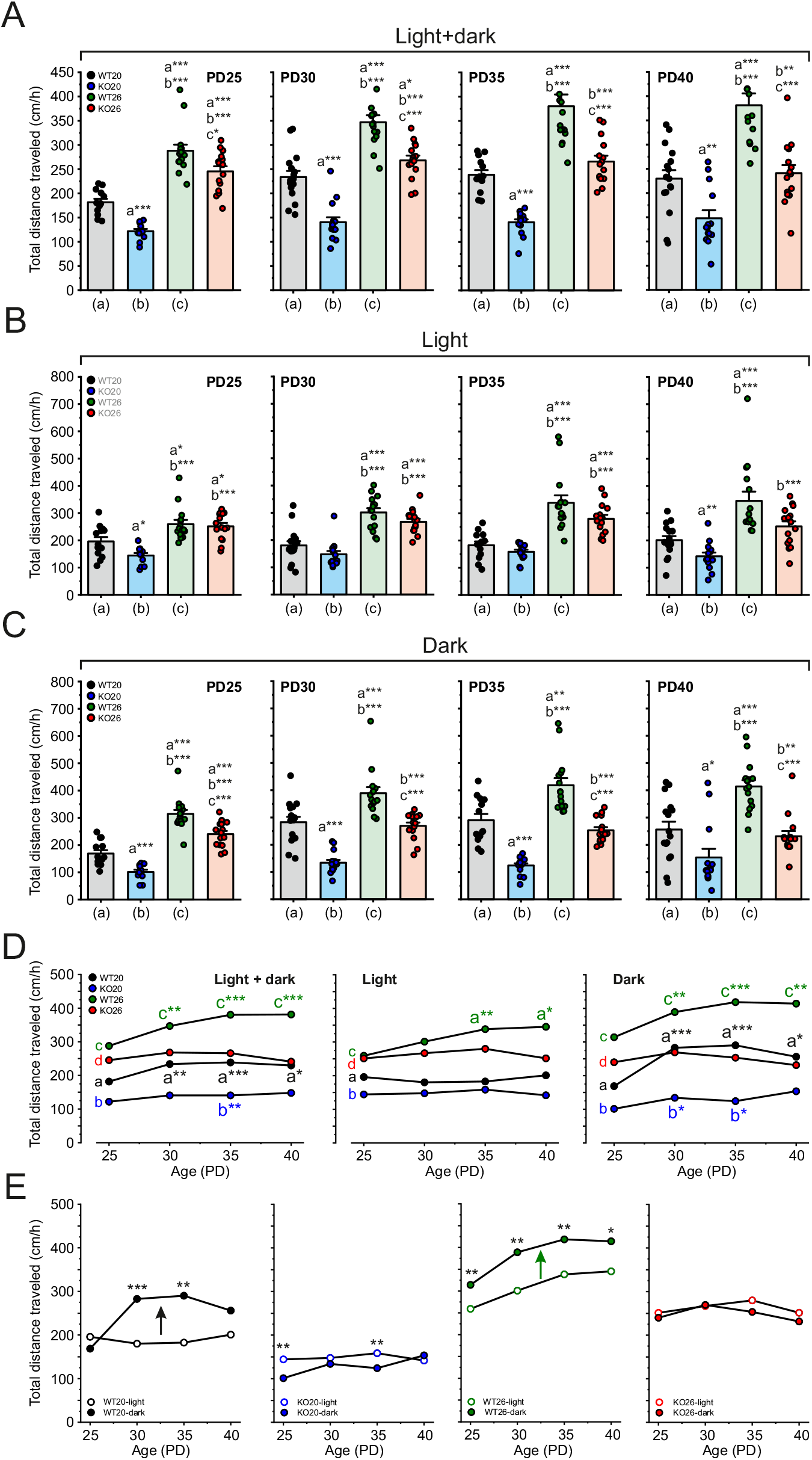
Effect of increased ambient temperature on the voluntary movement activity of WT and KO mice. (**A**) Total distance travelled (cm/h) during the total time period (light + dark phase) compared for WT20, KO20, WT26 and KO26 mice of different age (PD25, PD30, PD35, PD40). (**B**) Same as panel A, but now during light phase. (**C**) Same as panel A, but now during the dark phase. (**D**) Comparison of the total distance travelled (cm/h) for WT20, KO20, WT26 and KO26 mice as a function of age (PD) during 24 h, during the light phase, and during the dark phase. (**E**) Comparison of the total distance travelled (cm/h) between the light phase (open symbols) and dark phase (filled symbols) for WT20, KO20, WT26 and KO26 mice as a function of age (PD). **Statistics:** Datapoints in panel A, B and C reflect individual mice (n=12-16 mice for each of the four conditions). In panel A, B and C, data was compared within each PD between WT20, KO20, WT26 and KO26. The average data from panels A, B, C was used to generate panel D and E (error bars and individual data points were omitted for visualization purposes). In panel D, data at PD30, PD35 and PD40 was compared with PD25 (a,b,c,d). In panel E, data for WT20, KO20, WT26 and KO26 was compared between the light phase and dark phase at each PD. All significant differences between conditions (a,b,c,d) are marked by: *p<0.05, **p<0.01 or ***p<0.001.

### Effect of increased ambient temperature on skin and core temperature, body weight and -related parameters and food intake

Next, we determined how a higher T_A_ impacted on skin temperature (T_S_), core temperature (T_C_), BW and -related parameters and food intake (FI) in WT and KO mice, as schematically depicted (**Fig. 5A**). This also included analysis of whole-body energy metabolism by indirect calorimetry (InCa; **Fig. 6**). To retain sufficient mice for analysis, measurements were performed prior to occurrence of KO death (**Fig. 2B**). Average T_S_ values (**Fig. 5B**; bottom half) were (mean±SD; full age period): 29.9±0.7 °C (WT20), 29.8±0.8 °C (KO20), 32.3±0.8 °C (WT26) and 32.2±0.7 °C (KO26). These results demonstrate that T_S_ increases by ∼2.5 °C at the higher T_A_ (p<0.001). Average T_C_ values (**Fig. 5B**; top half) were: 36.1±0.583 °C (WT20), 36.0±0.725 °C (KO20), 37.5±0.3 °C (WT26) and 37.4±0.3 °C (KO26). This demonstrates that T_C_ increases by ∼1.5 °C at the higher T_A_ (p<0.001). The T_C_ of KO mice was apparently lower relative to WT20 mice after PD34 (**Fig. 5B**; boxes and p-values). In line the data presented above (**Fig. 2D-E**), KO20 and KO26 mice displayed an attenuated BW gain and reached their reduced maximum BW earlier than WT20 and WT26 mice (**Fig. 5C**). FI was identical for WT20 and KO20 mice, and similarly reduced for WT26 and KO26 mice (**Fig. 5D**). It was previously demonstrated that murine FI increases as a function of BW (**Bachmanov et al., 2002**). Since KO mice displayed a lower BW than WT mice (**Fig. 2D** and **Fig. 5C**), the total FI (energy) required per gram BW gain (FI/ΔBW) was higher in KO than in WT mice (**Fig. 5E**), indicating a lower food efficiency. Regarding BW gain, this was less for KO relative to WT mice, less for WT26 relative to WT20 mice, and identical for KO20 and KO26 mice (**Fig. 6F**). A similar pattern was observed for lean mass (LM) and fat mass (FM) gain, being reduced in KO relative to WT mice (**Fig. 6G-H**). We conclude that an elevated T_A_: (**1**) increases T_S_ and T_C_ to similar values in WT and KO mice, (**2**) reduces FI in WT and KO mice, (**3**) does not or minimally affect BW, LM and FM gain in WT and KO mice.

**Figure 5:**
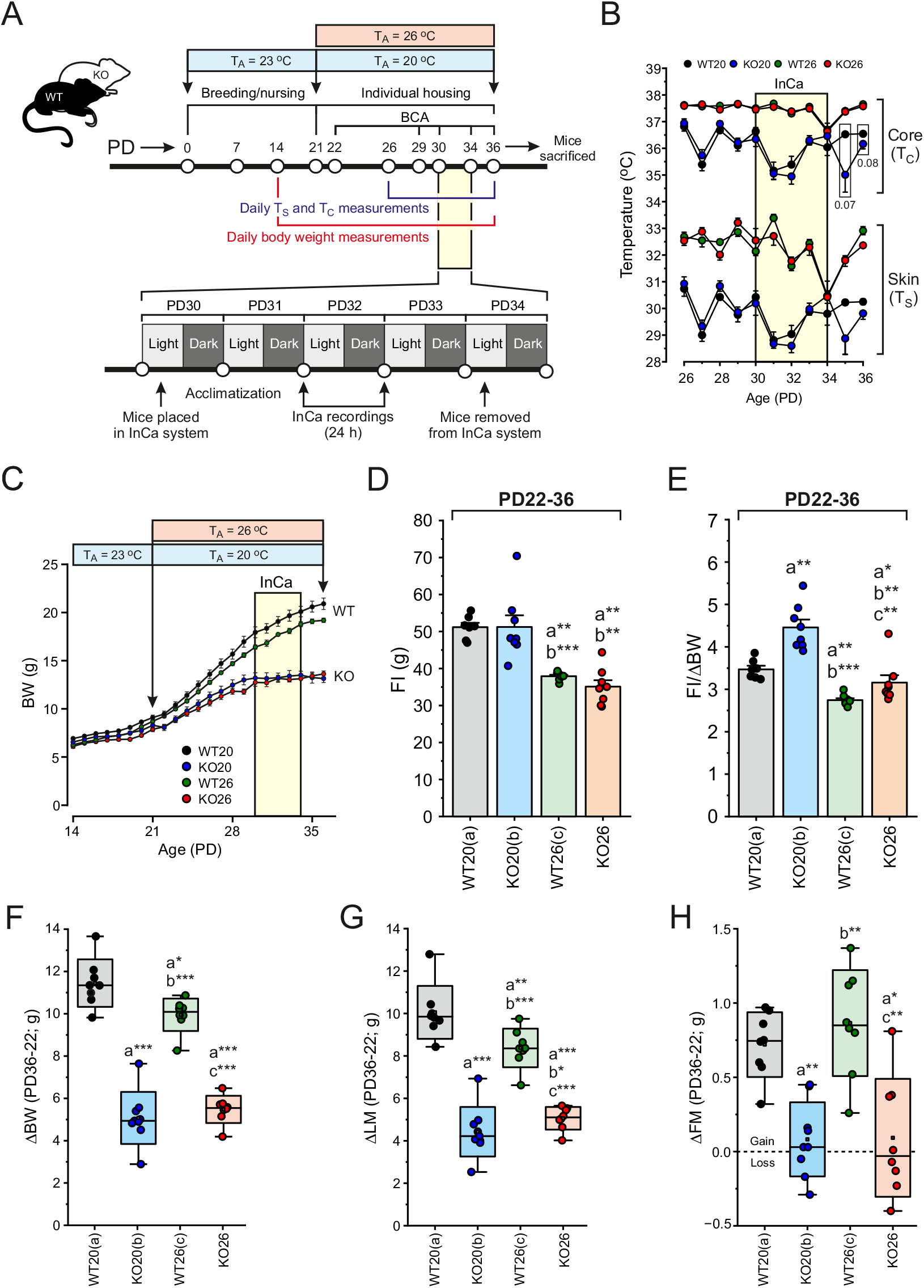
Effect of increased ambient temperature on skin temperature, core temperature, body weight, food intake and body composition of WT and KO mice. (**A**) Design of the indirect calorimetry (InCa) study. The time during which the mice were placed in the InCa system is highlighted in yellow. BCA = body composition analysis; PD = postnatal day; T_A_ = ambient temperature; T_C_ = core temperature; T_S_ = skin temperature. (**B**) T_C_ and T_S_ of WT and KO mice as a function of age, housed at a T_A_ of 20 °C (WT20, KO20) or 26 °C (WT26, KO26). The time during which the mice were placed in the InCa system is highlighted in yellow. (**C**) Body weight (BW) as a function of age for WT20, KO20, WT26 and KO26 mice. The time during which the mice were placed in the InCa system is highlighted in yellow. (**D**) Total food intake (FI) between PD22-36 for WT and KO mice housed at a T_A_ of 20 °C (WT20, KO20) or 26 °C (WT26, KO26). (**E**) Total FI corrected for BW increase (ΔBW) between PD22-36. (**F**) Change in BW (ΔBW) between PD22-36. (**G**) Change in lean mass (ΔLM) between PD22-36. (**H**) Change in fat mass (ΔFM) between PD22-36. **Statistics:** Data in panel B, C, D and E was obtained from n=8 mice for each condition. Data in panel F, G and H was obtained from n=8 mice (WT20, WT26, KO26) and n=9 mice (KO20). In panel F, G and H, error bars mark the 99% (upper) and 1% (lower) percentile, the boundary boxes mark the standard deviation (SD), the square marks the mean value of the data, and the horizontal line within the box indicates the median value of the data. Significant differences between the indicated conditions in panel B (boxes) and panel D, E, F, G and H (a,b,c) are marked by: *p<0.05, **p<0.01, ***p<0.001 or the exact p-value (borderline significance).

**Figure 6:**
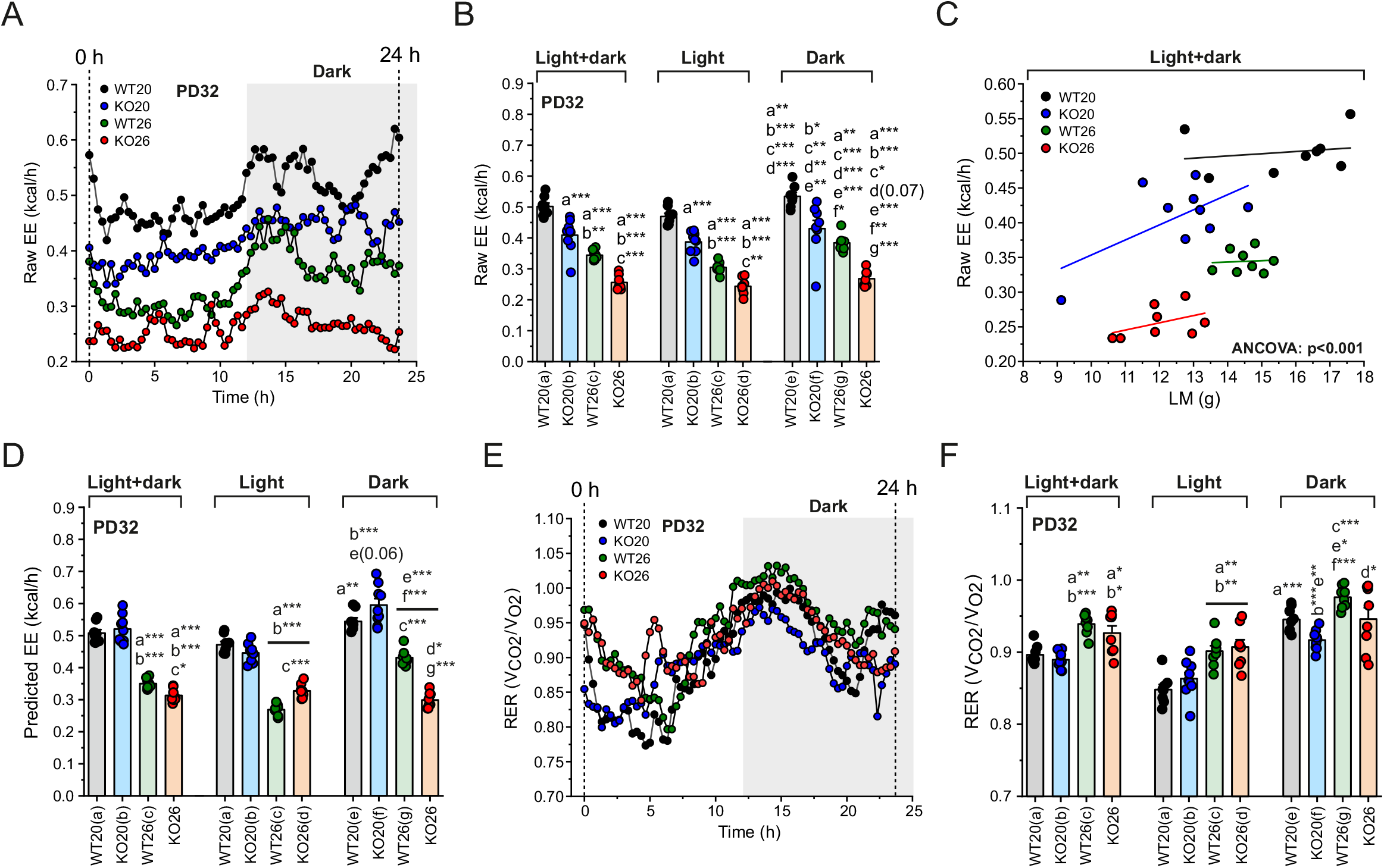
Effect of increased ambient temperature on energy expenditure and respiratory exchange ratio in WT and KO mice. (**A**) Average raw energy expenditure (EE) obtained by indirect calorimetry (InCa) during the total time period (light + dark phase) for WT and KO mice housed at a T_A_ of 20 °C (WT20, KO20) or 26 °C (WT26, KO26). (**B**) Raw EE total time period (light + dark phase), during the light phase, and during the dark phase. (**C**) Relationship between raw EE and lean mass (LM) for the four experimental groups during the total time period (light + dark phase), evaluated by analysis of covariance (ANCOVA). Lines represent linear fits to the data. (**D**) Predicted EE for WT and KO mice during the total time period (light + dark phase), during the light phase, and during the dark phase. (**E**) Same as panel A, but now for the average respiratory exchange ratio (RER). (**F**) RER values for WT and KO mice during the total time period (light + dark phase), during the light phase, and during the dark phase. **Statistics:** The data in this figure was obtained from n=8 mice (WT20, WT26, KO26) and n=9 mice (KO20). Data in panel B, D and F was compared within the 24 h period (a,b,c) and within and between the dark and light period (a,b,c,d,e,f,g). All significant differences between conditions (a,b,c,d,e,f,g) are marked by: *p<0.05, **p<0.01, ***p<0.001 and/or the exact p-value (borderline significance).

### Effect of increased ambient temperature on whole-body energy metabolism

Next, whole-body energy metabolism was analyzed for WT and KO mice using InCa (**Fig. 5A**). Data was analyzed for the total time period (24 h), light phase and dark phase To retain sufficient mice for analysis, they were studied prior to KO death at PD32 (**Fig. 2B**). Increasing T_A_ reduced the energy expenditure (EE) of WT and KO mice over the complete time period, as well as during the light phase and dark phase (**Fig. 6A-B**). Both WT and KO mice displayed a higher “raw” EE during the dark phase than during the light phase, but KO mice always displayed a lower raw EE relative to WT mice (**Fig. 6B**). Given the LM differences between WT and KO mice (**Fig. 5C**) the obtained raw EE data were normalized (**Fig. 6C**; **Tschöp et al., 2012; Müller et al., 2021**). These “predicted EE” values were also reduced at higher T_A_ for WT and KO mice (**Fig. 6D**). Over the complete time period, the predicted EE was similar for WT20 and KO20 mice, but lower for KO26 relative to WT26 mice (**Fig. 6D**). Separating the light and dark phase, revealed a higher predicted EE in the dark phase for WT20, KO20 and WT26, but not for KO26 (**Fig. 6D**). Moreover, while WT26 was lower than KO26 in the light phase, the opposite was the case in the dark phase (**Fig. 6D**). To gain insight into the relative substrate usage for energy production, the respiratory exchange ratio (RER) was analyzed (**Fig. 6E**). RER values routinely range between 0.70 (100% fat oxidation) and 1.00 (100% glucose oxidation), but can also exceed the latter value (*e*.*g*. in case of lipogenesis). Over the total time period and during the light phase, RER values were higher in WT and KO mice at higher T_A_, with no observed differences between WT and KO mice (**Fig. 6F**). During the dark phase, RER values were higher than during the light phase for WT and KO mice, suggesting a shift in metabolism from fat oxidation towards glucose oxidation. At both temperatures, RER values were apparently lower in KO mice than in WT mice during the dark phase (**Fig. 6F**). Taken together, the results in this section demonstrate that: (**1**) an elevated T_A_ reduces predicted EE and increases RER, and (**2**) predicted EE and RER are differentially affected during the light and dark period between WT26 and KO20 mice.

## DISCUSSION

*Ndufs4*^*-/-*^ mice are a widely used animal model for pediatric LS. Here we demonstrate that housing these mice at T_A_ = 26 °C (KO26 mice) instead of T_A_ = 20 °C (KO20 mice) reduces energy expenditure (EE), increases lifespan, reverses pathology in specific brain regions, and increases voluntary locomotor activity.

### Increased ambient temperature elevates skin, core and brain temperature in KO mice

Our findings provide evidence that KO26 mice display a higher T_S_, T_C_ and brain temperature than KO20 mice (**Fig. 2F-G**; **Fig. 5B**). Both T_S_ and T_C_ fluctuated with age (**Fig. 5B**). These fluctuations were apparently smaller at increased T_A_, and were previously linked to thermoregulation and variations in metabolism and other physiological processes (**Gordon, 2017; Škop et al., 2024**). Although detailed empirical studies are lacking, it appears that LS/LSS/MD patients also exhibit a lower brain temperature and/or abnormal thermoregulation (**Noorda et al., 2012; Rango et al., 2014; Sofou et al., 2014; Chang et al., 2020; Ball et al., 2024**).The latter might be linked to fever-induced hypothermia (**Yagasaki et al., 2019**) and infection-induced worsening of symptoms often observed in LS patients (**Baertling et al., 2014; Chang et al., 2020**). This is supported by the fact that thermoregulation during cold-adaptation and fever are mediated by similar mechanisms (**Guzmán-Ruiz et al., 2015; Nakamura, 2011; Tansey & Johnson, 2015**). In this context, the slower breathing rate observed in KO mice and LS patients with *NDUFS4* mutations (**Quintana et al., 2012**), might reflect an adaptive response to reduce heat loss (**Kozyreva, 2013**). With respect to thermoregulation, the hypothalamus acts as a coordinating center and contains temperature-sensitive neurons in the preoptic area (**Boulant, 2000; Tan & Knight, 2018; Tran et al., 2022**). Therefore, the observed hypothalamo-pituitary dysfunction in a patient with Mitochondrial Encephalopathy, Lactic Acidosis, and Stroke-like episodes (MELAS; **Joko et al., 1997**), might suggest hippocampal involvement in MD. Moreover, hypothalamus-mediated thermoregulation is linked to the storage of cold-sensitive memory engrams in the hippocampus (**Muñoz-Zamora et al., 2025**). This suggests that the normalization of hippocampal GFAP signals in KO26 mice (**Fig. 3A-B**) might reflect an (indirect) effect on thermoregulatory pathways.

### Increased ambient temperature reverses astrocyte activation-linked brain pathology in the hippocampus and motor cortex of KO mice

Astrocyte activation (GFAP-positive cells) was increased in the hippocampus, olfactory bulb and motor cortex of KO20 mice (**Fig. 3**). Remarkably, this activation was reduced in the hippocampus and motor cortex, but not in the olfactory bulb, of KO26 mice. This strongly suggests that astrocyte activation in the hippocampus and motor cortex of KO20 mice is not a direct consequence of *Ndufs4* gene deletion, but result from a too low brain temperature. Conversely, astrocyte activation in the olfactory bulb was not mitigated in KO26 mice, strongly suggesting that this activation is a true pathological consequence of *Ndufs4* gene deletion. Microglial activation was demonstrated in post-mortem brain samples from LS patients (**Daneshgar et al., 2023**). With respect to microglial activation (IBA1-positive cells), we did not observe a clear pattern of differences between KO20 and KO26 mice (**Fig. 3**). This suggests that altered microglial activation in the analyzed brain regions is not linked to disease mitigation in KO26 mice. It was previously demonstrated that the total number of changed proteins (KO20/WT20) was lower in the hippocampus (70 proteins) than in the superior colliculus (101 proteins; **van de Wal et al., 2025**). Analysis of these changes suggested a pro-pathological upregulation of fatty acid oxidation (FAO) and synthesis (FAS) pathways in the hippocampus but not in the superior colliculus (**van de Wal et al., 2025**). Adaptive upregulation of FAS pathways was also demonstrated in the olfactory bulb, but not in the cerebellum and brainstem of KO20 mice (**Khumalo et al., 2025**). In the context of the current study, these findings suggest that the hippocampal and olfactory bulb regions are unable to adapt to the *Ndufs4* gene knockout in KO20 mice (displaying FAO/FAS upregulation and increased astrocyte activation). In contrast, the superior colliculus region displayed a successful adaptation to *Ndufs4* gene knockout in KO20 mice (no FAO/FAS upregulation and no increased astrocyte activation).

### Increased ambient temperature stimulates voluntary activity

Focusing on the motor cortex, we demonstrated that increased astrocyte activation in KO20 mice was reduced in KO26 mice (**Fig. 3E-F**). Because this reduction was paralleled by increased voluntary movement activity in KO26 relative to KO20 mice (**Fig. 4A-B-C**), we conclude that this GFAP-linked pathology reversal is functionally relevant. WT20 were also more active than WT26 mice, compatible with the fact that healthy mice run a greater distance on a wheel when housed at thermoneutrality relative to room temperature (**Lac et al., 2023**). KO26 mice were always more active than KO20 mice. However, in contrast to WT mice, KO mouse activity neither increased with age (**Fig. 4D**), nor during the dark phase (**Fig. 4E**). This demonstrates that although the activity of KO26 mice is increased, it is not restored to WT26 levels. We speculate that an energetic limitation or preferential demand of other processes may prevent full restoration. Similar to KO20 mice, pediatric MD patients displayed a significantly lower peak activity and were resting more, compared to their age- and sex-matched peers (**Koene et al., 2017**).

### Increased ambient temperature reduces food intake, lowers energy expenditure, and increases respiratory exchange ratio

KO mice displayed a lower body weight (BW), BW gain (ΔBW), lean mass gain (ΔLM) and fat mass gain (ΔFM) relative to WT mice and this difference was not affected by increased T_A_ (**Fig. 2D** and **Fig. 5C-F-G-H**). In contrast, food intake (FI) and the FI per gram BW gain (FI/ΔBW) were reduced at higher T_A_ for WT and KO mice (**Fig. 5D-E**), Comparison of WT and KO mice revealed that FI/ΔBW was higher in KO than in WT mice (**Fig. 5E**). This means that KO mice require a higher energy intake relative to WT mice to gain a similar amount of BW (**Fig. 5D**), indicating an apparent lower food conversion rate. This is supported by the fact that KO26 mice do not gain fat mass (**Fig. 5H**). Moreover, KO mice displayed no EE increase in the dark phase (**Fig. 6B-D**). Similar to previous studies in healthy mice, an increased T_A_ lowered EE (**Fig. 6A-D**) because less energy is required to maintain T_B_ (**Fischer et al., 2016; John et al., 2022; Sadler et al, 2022**). Elevating T_A_ also reduced EE in healthy human individuals (**Henkel et al., 2025**). Interestingly, it was previously demonstrated that thermal insulation in mice was not increased by obesity of any kind (**Fischer et al., 2016**). This suggests that heat loss is not enhanced in KO mice due to their lower FM. Related to this, *Ndufs4*^*-/-*^ mice partially lose their hair, which grows back during the next hair growth cycle (**Kruse et al., 2008**). Although not systematically documented, we also visually observed this temporary hair loss during our InCa experiments. Potentially, hair loss could enhance heat loss, thereby increasing EE in KO mice (**Fisher et al., 2016**). However, the latter study reported that although EE was higher in shaved mice than in control mice at T_A_ = 20°C, these mice displayed a virtually identical EE at T_A_ = 26 °C. Therefore, it is expected that KO20 mice exhibit a higher EE than WT20 mice, whereas KO26 and KO26 mice display a similar EE. Because this pattern was not observed (**Fig. 6B-D**), we conclude that temporarily hair loss is not primarily responsible for the EE differences between WT and KO mice in the current study. Pediatric MD patients displayed a borderline significant increase in resting energy expenditure (REE) relative to controls (p=0.086). In contrast, analysis of MD patients carrying one of the most common pathogenic mtDNA variants (m3242A>G; **Li et al., 2022**) revealed a borderline-significant decrease in REE (p=0.052) relative to controls (**Zweers et al., 2021**). In yet another study, no differences in REE between pediatric MD patients and healthy controls were observed (**Fiuza-Luces et al., 2016**). We conclude that more empirical studies are required to determine if and how EE is affected in LS and other MDs.

### Mechanistic aspects

MD patients exhibit a significantly reduced BW, FM and body mass index (BMI; **de Laat et al., 2015; Hou et al., 2019**). This suggests that, similar to KO20 mice (**Chen et al., 2022**), MD patients maintain their T_B_ by increasing FAO, potentially reducing their energy reserves (*i*.*e*. adipose tissue) or paralleled by a decreased food efficiency (*i*.*e*. the gram weight gain per unit of energy intake). Related to this, MD patients frequently suffer from malnutrition and might benefit from nutritional interventions (**DiVito et al., 2023**). Mechanistically, T_B_ changes can functionally alter the properties of proteins, nucleic acids, membranes and the physicochemical properties of the cytosol and intra-organelle solvents. These alterations include protein conformational changes, protein aggregation, base-pairing stability, membrane fluidity, transcriptome alterations and biomolecule reaction and diffusion (**Bulthuis et al., 2023; Shin et al., 2024; Conti & de Cabo, 2025**). Analysis of isolated mouse cortical mitochondria revealed that mitochondrial oxygen consumption was lower and reactive oxygen species (ROS) production was higher at 32 °C relative to 37 °C (**Ali et al., 2010**). Similarly, mitochondria in permeabilized mouse brain displayed lower respiration and OXPHOS maximal rates at 28 °C relative to 37 °C (**Pamenter et al., 2018**). This suggests that the lower T_B_ and brain temperature in KO20 mice further aggravate the already reduced mitochondrial activity reported in *Ndufs4*^*-/-*^ mice (**Kruse et al., 2008**), supporting a mechanism in which mitochondrial oxygen consumption and ATP production is higher in KO26 than in KO20 mice. Such a mechanism could also increase mitochondrial oxygen consumption in KO26 mouse brain, thereby possibly mitigating its hyperoxic state leading to lifespan extension (**Jain et al., 2016; Blume et al., 2025**). The fact that a lower T_B_ is not associated with pathology in WT20 mice supports the conclusion that a reduced CI activity is the primary disease driver in *Ndufs4*^*-/-*^ mice. This is compatible with recent evidence that neuroinflammatory responses are secondary to mitochondrial deficiency (**Kozicz & Morava, 2025**). Expression of brain OXPHOS subunits was similar in KO20 and KO26 mice (**Fig. 2H-I**), arguing against OXPHOS activity being stimulated in the latter mice.

### Conclusion

We demonstrate that specific brain and -related aberrations in *Ndufs4*^*-/-*^ mice are reversed or mitigated by increasing T_A_. This suggests that these aberrations are not primary consequences of *Ndufs4* gene deletion and ensuing mitochondrial dysfunction. Our results advocate the use of (more) thermoneutral housing to evaluate pathomechanisms and intervention strategies in murine models of human disease. Given their reduced mitochondrial energy production and aberrant thermoregulation, lowering energy requirements in LS/LSS and other MD patients might have therapeutic value and/or improve their quality of life.

## Potential conflict of interest

Christian J.M.I. Klein is employed at TSE Systems GmbH (Berlin, Germany). However, this company had no involvement in the data collection, analysis and interpretation, writing of the manuscript, nor in the decision to submit the manuscript for publication.

## Data availability

All data is included with this work or available from the authors at reasonable request.

## Acknowledgements including funding

- Melissa A.E. van de Wal was funded by a Radboudumc Junior Researcher Grant awarded to Clara van Karnebeek and Werner J.H. Koopman.
- Christian J.M.I. Klein is an ESR fellow of the INSPIRE European Training Network. INSPIRE received funding from the EU Horizon 2020 Research and Innovation program, under the Marie Skłodowska-Curie program Grant (GA858070)
- Werner J.H. Koopman and Judith Homberg were supported by Radboudumc Research Group Leader (RGL) funding.
- Indirect calorimetry (InCa) experiments were supported by a United for Metabolic Diseases Catalyst Grant to Melissa A.E. van de Wal and Werner J.H. Koopman (UMD-CG-2022-001).
- We thank A. van der Ven (Department of Medical BioSciences, Radboud University Medical Center, Nijmegen, The Netherlands) for assistance with OXPHOS Western blotting.
- We thank J. de Deugd (Human and Animal Physiology, Wageningen University, Wageningen, The Netherlands) for assistance with the InCa experiments.

## Author contributions

- Melissa A.E. van de Wal = manuscript writing, Western blotting, brain pathology analysis, mouse voluntary movement experiments, data analysis, discussions, funding
- Christian J.M.I. Klein = manuscript writing, InCa experiments, data analysis/interpretation, discussions
- Merel J.W. Adjobo-Hermans = manuscript writing, discussions, co-supervision
- Els van de Westerlo = technical support, manuscript writing
- Clara van Karnebeek = manuscript writing, clinical aspects, co-supervision, funding
- Mirian Janssen = manuscript writing, clinical aspects, discussions
- Jan C. van der Meijden = manuscript writing, clinical aspects, discussions
- Judith R. Homberg = manuscript writing, data interpretation, supervision of voluntary movement studies
- Jaap Keijer = manuscript writing, data interpretation, discussions
- Evert M. van Schothorst = manuscript writing, supervision of InCa experiments, data analysis/interpretation, discussions
- Werner J.H. Koopman = manuscript writing, data analysis/interpretation, overall supervision, funding

## Abbreviations

AOD: age of death
BCA: body composition analysis
BW: body weight
CI: complex I
CIRBP: cold-inducible RNA-binding protein
CT: control
EE: energy expenditure
FI: food intake
FM: fat mass
G3P: glycerol-3-phosphate
GFAP: glial fibrillary acidic protein
IBA1: ionized calcium-binding adapter molecule 1
InCa: indirect calorimetry
IR: infra-red
KO: whole-body *Ndufs4* knock-out
KO20: KO mice housed at an ambient temperature of 20 °C
KO26: KO mice housed at an ambient temperature of 26 °C
LS: Leigh syndrome
LLS: Leigh-like syndrome
LM: lean mass
LSS: Leigh syndrome spectrum
MD: mitochondrial disease
mtDNA: mitochondrial DNA
NA: nicotinic acid
nDNA: nuclear DNA
NDUFS4: NADH dehydrogenase [ubiquinone] iron-sulfur protein 4
OXPHOS: oxidative phosphorylation
PD: postnatal day
RBM3: RNA-binding motif 3
RER: respiratory exchange ratio
T_A_: ambient temperature
T_B_: body temperature
T_C_: core temperature
T_S_: skin temperature
WT: wildtype
WT20: WT mice housed at an ambient temperature of 20 °C
WT26: WT mice housed at an ambient temperature of 26 °C

---

**Supplementary Figure S1:**
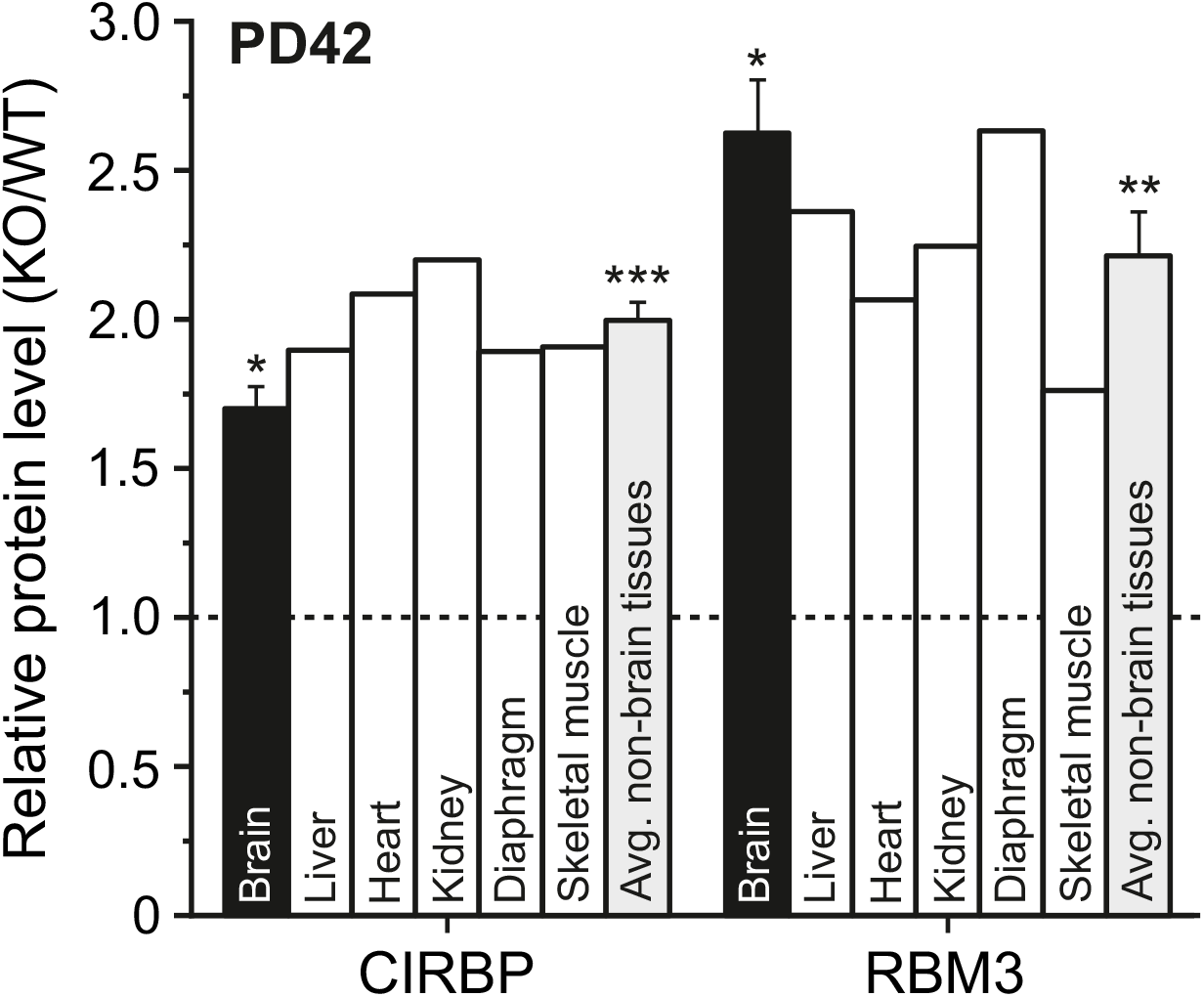
Expression of cold-induced proteins in KO mice. Relative protein level (KO/WT) of the cold-inducible proteins CIRBP and RBM3 in brain (black bars) and other tissues (open bars). The gray bar represents the average protein level (KO/WT) in the non-brain tissues (data obtained by proteome analysis; see **Supplementary Table S3**).

**Supplementary Figure S2:**
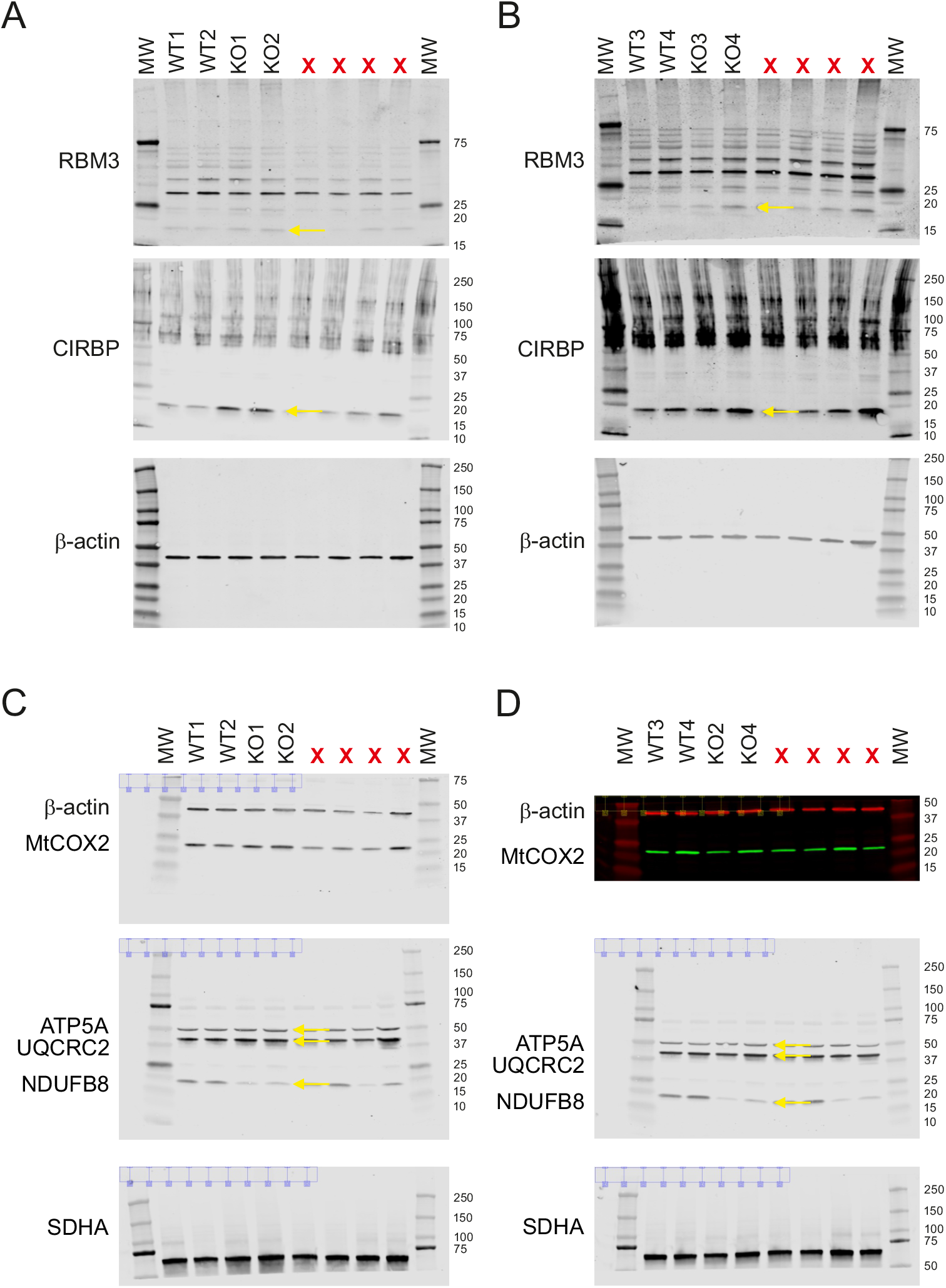
Original Western blots used for Figure 2F-G-H-I. The data in panel A and C were used for **Fig. 2F** and **H**. All data was quantified to generate **Fig. 2G** and **I**. (**A**) Western blot analysis of RBM3 (arrow), CIRBP (arrow) and β-actin (normalization) in two brain samples from wildtype (WT) mice (WT1, WT2), and two brain samples of *Ndufs4*^*-/-*^ knockout (KO) mice (KO1, KO2). MW indicates molecular weight markers in kDa (numbers): RBM3 (predicted MW = 16.6-kDa), CIRBP (18.6-kDa) and β-actin (42-kDa). Lanes marked with “X” were not part of the current study. (**B**) Same as panel A, but now for two additional WT (WT3, WT4) and KO (KO3, KO4) samples. (**C**) Same as panel A, but now for NDUFB8 (CI; 18-kDa; arrow), SDHA (CII; 72-kDa), UQRCRC2 (CIII; 48-kDa; arrow), MtCOX2 (CIV; 26-kDa), ATP5A (CV; 60-kDa; arrow) and β-actin (normalization; 42-kDa). (**D**) Same as panel C, but now for two additional WT (WT3, WT4) and KO (KO3, KO4) samples.

**Supplementary Table S1:**
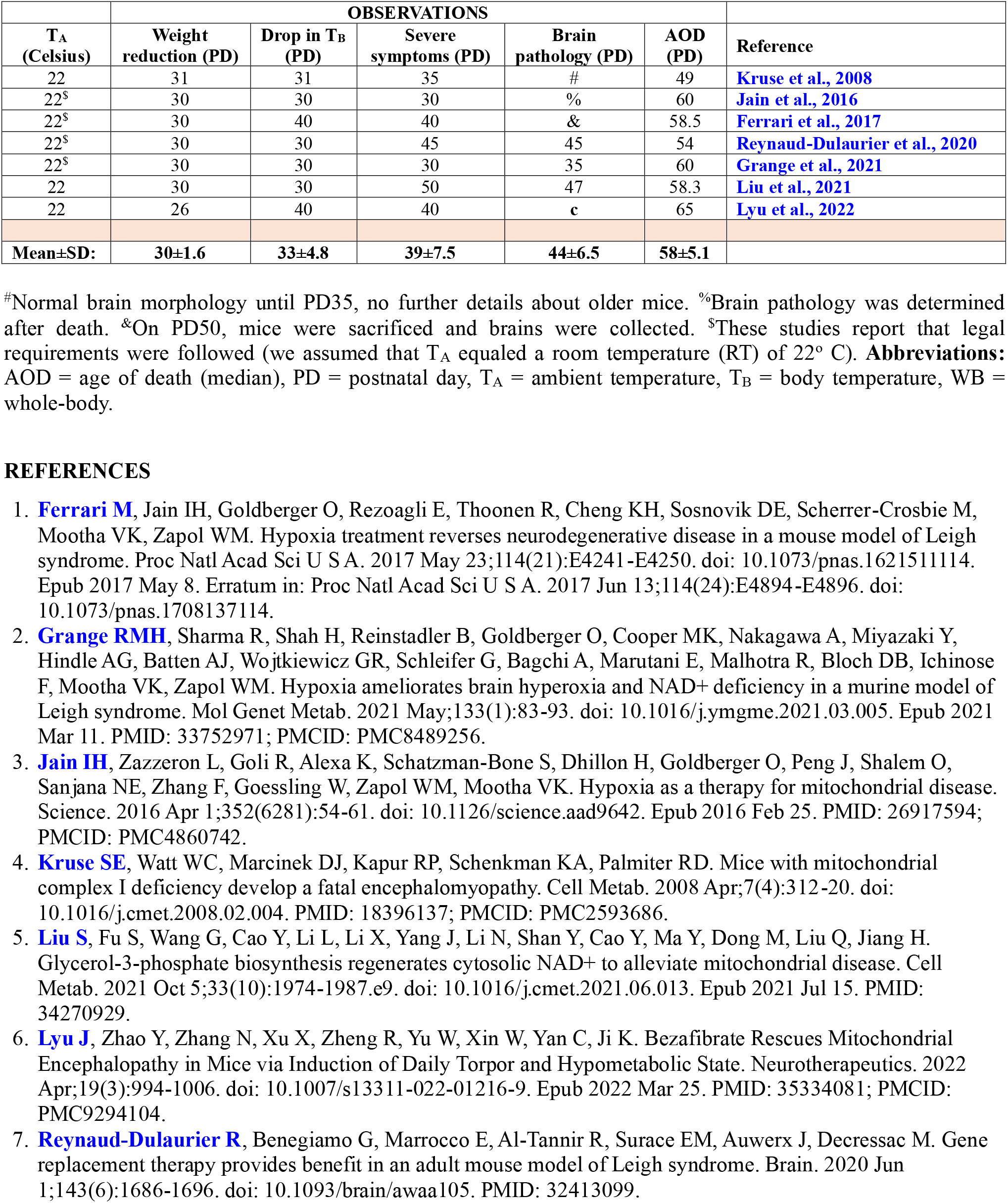
Phenotype in WB-KO mice over time.

**Supplementary Table S2:** Body temperature (T_B_) and age of death (AOD) of untreated and treated WB-KO mice.

| Untreated |  | Treated |  |  | Reference |
| --- | --- | --- | --- | --- | --- |
| T<sub>B</sub> | AOD | Treatment | T<sub>B</sub> | AOD |  |
| 35.0 | 65 | Vehicle | -- | -- | Blume et al., 2025 |
| -- | -- | HypoxyStat | 36.5 | 180 | Blume et al., 2025 |
| -- | -- | GBT-440 | 36.2 | 90 | Blume et al., 2025 |
| 32.5 | 58.3 | Increasing G3P | 36 (PD50) | 84.0 | Liu et al., 2021 |
| 33.0 | 72 (17% O<sub>2</sub>) | NA +17% O<sub>2</sub> | 34.5 | 104 | Grange et al., 2021 |
| 34.0 | 54 | AAV.PHP.B-*hNDUFS4* | 37.0 | 250 | Reynaud-Dulaurier et al., 2020 |
| 33.0 | 60 | Hypoxia (11% O<sub>2</sub>) | 37.0<sup>#</sup> | 170 | Jain et al., 2016 |
| 33.0 | 49 | No treatment | -- | -- | Kruse et al., 2008 |

**Abbreviations:** AAV.PHP.B-*hNDUFS4* = adeno-associated virus (AAV) delivered human *NDUFS4* (*hNDUFS4*); AOD = age of death (in postnatal days; PD); G3P = glycerol-3-phosphate; NA = nicotinic acid; T<sub>B</sub> = body temperature in degrees Celsius; # = reported as “no drop” in T<sub>B</sub>.

**Supplementary Table S3:** Expression of cold-induced proteins in WB-KO mouse tissues.

| Name | UniProt ID | Fold change in protein expression (KO/WT) |  |  |  |  |  | Mean non-brain tissues (±SEM) |
| --- | --- | --- | --- | --- | --- | --- | --- | --- |
|  |  | Mean brain (±SEM) | Liver | Heart | Kidney | Diaphragm | Skeletal muscle |  |
| CIRBP | P60824 | 1.70 ±0.0750<br>BH-P<0.05 (*) | 1.90 | 2.09 | 2.20 | 1.89 | 1.91 | 2.00±0.0625<br>p<0.001 (***) |
| RBM3 | O89086 | 2.63 ±0.184<br>BH-P<0.05 (*) | 2.63 | 2.07 | 2.25 | 2.63 | 1.76 | 2.21±0.145<br>P<0.01 (**) |

For the brain three WT and three KO animals were analyzed (brain slices). For all other tissues one WT animal and one KO animal were analyzed. Details on the followed proteomic strategy are provided elsewhere (Adjobo-Hermans et al., 2020 ; van de Wal et al., 2025). BH-P = Benjamini-Hochberg p-value. \*\* = p<0.01 (one-population Student’s t-test against 1.0), \*\*\* = p<0.001 (one-population Student’s t-test against 1.0). SEM = standard error of the mean. In case of the brain data, the standard deviation (SD<sub>WT/KO</sub>) of the fold change in protein expression (WT/KO) was determined directly from the normalized relative abundance (RA) of CIRBP and RBM3 in the samples (3 WT and 3 KO) for CIRBP (WT) = 13.2, 12.4, 11.4; CIRBP (KO) = 21.1, 20.5, 21.4; RBM3 (WT) = 9.36, 10.1, 8.11 and RBM3 (KO) = 25.6, 23.7, 23.2 (data taken from Van de Wal et al., 2025). The SD<sub>KO/WT</sub> for CIRBP and RBM3 was calculated using:

## REFERENCES

1. Abreu-Vieira G, Xiao C, Gavrilova O, Reitman ML. Integration of body temperature into the analysis of energy expenditure in the mouse. Mol Metab. 2015 Mar 10;4(6):461–70. doi: 10.1016/j.molmet.2015.03.001. PMID: 26042200; PMCID: PMC4443293.

2. Adjobo-Hermans MJW, de Haas R, Willems PHGM, Wojtala A, van Emst-de Vries SE, Wagenaars JA, van den Brand M, Rodenburg RJ, Smeitink JAM, Nijtmans LG, Sazanov LA, Wieckowski MR, Koopman WJH. NDUFS4 deletion triggers loss of NDUFA12 in Ndufs4-/-mice and Leigh syndrome patients: A stabilizing role for NDUFAF2. Biochim Biophys Acta Bioenerg. 2020 Aug 1;1861(8):148213. doi: 10.1016/j.bbabio.2020.148213. Epub 2020 Apr 23. PMID: 32335026.

3. Aguilar K, Comes G, Canal C, Quintana A, Sanz E, Hidalgo J. Microglial response promotes neurodegeneration in the Ndufs4 KO mouse model of Leigh syndrome. Glia. 2022 Nov;70(11):2032–2044. doi: 10.1002/glia.24234. Epub 2022 Jun 30. PMID: 35770802; PMCID: PMC9544686.

4. Ali SS, Marcondes MC, Bajova H, Dugan LL, Conti B. Metabolic depression and increased reactive oxygen species production by isolated mitochondria at moderately lower temperatures. J Biol Chem. 2010 Oct 15;285(42):32522–8. doi: 10.1074/jbc.M110.155432. Epub 2010 Aug 17. PMID: 20716522; PMCID: PMC2952254.

5. Anderson SL, Chung WK, Frezzo J, Papp JC, Ekstein J, DiMauro S, Rubin BY. A novel mutation in NDUFS4 causes Leigh syndrome in an Ashkenazi Jewish family. J Inherit Metab Dis. 2008 Dec;31 Suppl 2:S461–7. doi: 10.1007/s10545-008-1049-9. Epub 2008 Dec 26. PMID: 19107570.

6. Anglin RE, Rosebush PI, Noseworthy MD, Tarnopolsky M, Mazurek MF. Psychiatric symptoms correlate with metabolic indices in the hippocampus and cingulate in patients with mitochondrial disorders. Transl Psychiatry. 2012 Nov 13;2(11):e187. doi: 10.1038/tp.2012.107. PMID: 23149451; PMCID: PMC3565764.

7. Assereto S, Robbiano A, Di Rocco M, Rossi A, Cassandrini D, Panicucci C, Brigati G, Biancheri R, Bruno C, Minetti C, Trucks H, Sander T, Zara F, Gazzerro E. Functional characterization of the c.462delA mutation in the NDUFS4 subunit gene of mitochondrial complex I. Clin Genet. 2014 Jul;86(1):99–101. doi: 10.1111/cge.12248. Epub 2013 Sep 11. PMID: 24020637.

8. Assouline Z, Jambou M, Rio M, Bole-Feysot C, de Lonlay P, Barnerias C, Desguerre I, Bonnemains C, Guillermet C, Steffann J, Munnich A, Bonnefont JP, Rötig A, Lebre AS. A constant and similar assembly defect of mitochondrial respiratory chain complex I allows rapid identification of NDUFS4 mutations in patients with Leigh syndrome. Biochim Biophys Acta. 2012 Jun;1822(6):1062–9. doi: 10.1016/j.bbadis.2012.01.013. Epub 2012 Feb 3. PMID: 22326555.

9. Bachmanov AA, Reed DR, Beauchamp GK, Tordoff MG. Food intake, water intake, and drinking spout side preference of 28 mouse strains. Behav Genet. 2002 Nov;32(6):435–43. doi: 10.1023/a:1020884312053. PMID: 12467341; PMCID: PMC1397713.

10. Baertling F, Rodenburg RJ, Schaper J, Smeitink JA, Koopman WJ, Mayatepek E, Morava E, Distelmaier F. A guide to diagnosis and treatment of Leigh syndrome. J Neurol Neurosurg Psychiatry. 2014 Mar;85(3):257–65. doi: 10.1136/jnnp-2012-304426. Epub 2013 Jun 14. PMID: 23772060.

11. Ball M, Thorburn DR, Rahman S. Mitochondrial DNA-Associated Leigh Syndrome Spectrum. 2003 Oct 30 [updated 2024 May 9]. In: Adam MP, Feldman J, Mirzaa GM, Pagon RA, Wallace SE, Amemiya A, editors. GeneReviews [Internet]. Seattle (WA): University of Washington, Seattle; 1993–2024. PMID: 20301352.

12. Blume SY, Garg A, Martí-Mateos Y, Midha AD, Chew BTL, Lin B, Yu C, Dick R, Lee PS, Situ E, Sarwaikar R, Green E, Ramanan V, Grotenbreg G, Hoek M, Sinz C, Jain IH. HypoxyStat, a small-molecule form of hypoxia therapy that increases oxygen-hemoglobin affinity. Cell. 2025 Mar 20;188(6):1580-1588.e11. doi: 10.1016/j.cell.2025.01.029. Epub 2025 Feb 17. PMID: 39965572.

13. Bolea I, Gella A, Sanz E, Prada-Dacasa P, Menardy F, Bard AM, Machuca-Márquez P, Eraso-Pichot A, Mòdol-Caballero G, Navarro X, Kalume F, Quintana A. Defined neuronal populations drive fatal phenotype in a mouse model of Leigh syndrome. Elife. 2019 Aug 12;8:e47163. doi: 10.7554/eLife.47163. PMID: 31403401; PMCID: PMC6731060.

14. Boulant JA. Role of the preoptic-anterior hypothalamus in thermoregulation and fever. Clin Infect Dis. 2000 Oct;31 Suppl 5:S157–61. doi: 10.1086/317521. PMID: 11113018.

15. Bulthuis EP, Dieteren CEJ, Bergmans J, Berkhout J, Wagenaars JA, van de Westerlo EMA, Podhumljak E, Hink MA, Hesp LFB, Rosa HS, Malik AN, Lindert MK, Willems PHGM, Gardeniers HJGE, den Otter WK, Adjobo-Hermans MJW, Koopman WJH. Stress-dependent macromolecular crowding in the mitochondrial matrix. EMBO J. 2023 Apr 3;42(7):e108533. doi: 10.15252/embj.2021108533. Epub 2023 Feb 24. PMID: 36825437; PMCID: PMC10068333.

16. Budde SM, van den Heuvel LP, Smeets RJ, Skladal D, Mayr JA, Boelen C, Petruzzella V, Papa S, Smeitink JA. Clinical heterogeneity in patients with mutations in the NDUFS4 gene of mitochondrial complex I. J Inherit Metab Dis. 2003;26(8):813–5. doi: 10.1023/b:boli.0000010003.14113.af. PMID: 14765537.

17. Chang X, Wu Y, Zhou J, Meng H, Zhang W, Guo J. A meta-analysis and systematic review of Leigh syndrome: clinical manifestations, respiratory chain enzyme complex deficiency, and gene mutations. Medicine (Baltimore). 2020 Jan;99(5):e18634. doi: 10.1097/MD.0000000000018634. PMID: 32000367; PMCID: PMC7004636.

18. Chen X, Bollinger E, Cunio T, Damilano F, Stansfield JC, Pinkus CA, Kreuser S, Hirenallur-Shanthappa D, Roth Flach RJ. An assessment of thermoneutral housing conditions on murine cardiometabolic function. Am J Physiol Heart Circ Physiol. 2022 Feb 1;322(2):H234–H245. doi: 10.1152/ajpheart.00461.2021. Epub 2021 Dec 17. PMID: 34919456.

19. Conti B, de Cabo R. Promoting health and survival through lowered body temperature. Nat Aging. 2025 Apr 9. doi: 10.1038/s43587-025-00850-0. Epub ahead of print. PMID: 40205073.

20. Corrà S, Cerutti R, Balmaceda V, Viscomi C, Zeviani M. Double administration of self-complementary AAV9NDUFS4 prevents Leigh disease in Ndufs4-/-mice. Brain. 2022 Oct 21;145(10):3405–3414. doi: 10.1093/brain/awac182. PMID: 36270002; PMCID: PMC9586549.

21. Daneshgar N, Leidinger MR, L. S, Hefti M, Prigione A, Dai DF. Activated microglia and neuroinflammation as a pathogenic mechanism in Leigh syndrome. Front Neurosci. 2023 Jan 18;16:1068498. doi: 10.3389/fnins.2022.1068498. PMID: 36741056; PMCID: PMC9889986.

22. Davis MS, Bayly WM, Hansen CM, Barrett MR, Blake CA. Effects of hyperthermia and acidosis on mitochondrial production of reactive oxygen species. Am J Physiol Regul Integr Comp Physiol. 2023 Dec 1;325(6):R725–R734. doi: 10.1152/ajpregu.00177.2023. Epub 2023 Oct 9. PMID: 37811714.

23. de Groof AJ, Fransen JA, Errington RJ, Willems PH, Wieringa B, Koopman WJH. The creatine kinase system is essential for optimal refill of the sarcoplasmic reticulum Ca2+ store in skeletal muscle. J Biol Chem. 2002 Feb 15;277(7):5275–84. doi: 10.1074/jbc.M108157200. Epub 2001 Dec 4. PMID: 11734556.

24. de Haas R, Russel FG, Smeitink JA. Gait analysis in a mouse model resembling Leigh disease. Behav Brain Res. 2016 Jan 1;296:191–198. doi: 10.1016/j.bbr.2015.09.006. Epub 2015 Sep 9. PMID: 26363424.

25. de Haas R, Das D, Garanto A, Renkema HG, Greupink R, van den Broek P, Pertijs J, Collin RWJ, Willems PHGM, Beyrath J, Heerschap A, Russel FG, Smeitink JAM. Therapeutic effects of the mitochondrial ROS-redox modulator KH176 in a mammalian model of Leigh Disease. Sci Rep. 2017 Sep 15;7(1):11733. doi: 10.1038/s41598-017-09417-5. PMID: 28916769; PMCID: PMC5601915.

26. de Laat P, Zweers HE, Knuijt S, Smeitink JA, Wanten GJ, Janssen MC. Dysphagia, malnutrition and gastrointestinal problems in patients with mitochondrial disease caused by the m3243A>G mutation. Neth J Med. 2015 Jan;73(1):30–6. PMID: 26219939.

27. de Visser L, van den Bos R, Kuurman WW, Kas MJ, Spruijt BM. Novel approach to the behavioural characterization of inbred mice: automated home cage observations. Genes Brain Behav. 2006 Aug;5(6):458–66. doi: 10.1111/j.1601-183X.2005.00181.x. PMID: 16923150.

28. DiVito D, Wellik A, Burfield J, Peterson J, Flickinger J, Tindall A, Albanowski K, Vishnubhatt S, MacMullen L, Martin I, Muraresku C, McCormick E, George-Sankoh I, McCormack S, Goldstein A, Ganetzky R, Yudkoff M, Xiao R, Falk MJ R, Mascarenhas M, Zolkipli-Cunningham Z. Optimized Nutrition in Mitochondrial Disease Correlates to Improved Muscle Fatigue, Strength, and Quality of Life. Neurotherapeutics. 2023 Oct;20(6):1723–1745. doi: 10.1007/s13311-023-01418-9. Epub 2023 Sep 18. PMID: 37723406; PMCID: PMC10684455.

29. Fernández-Calleja JMS, Konstanti P, Swarts HJM, Bouwman LMS, Garcia-Campayo V, Billecke N, Oosting A, Smidt H, Keijer J, van Schothorst EM. Non-invasive continuous real-time in vivo analysis of microbial hydrogen production shows adaptation to fermentable carbohydrates in mice. Sci Rep. 2018 Oct 18;8(1):15351. doi: 10.1038/s41598-018-33619-0. PMID: 30337551; PMCID: PMC6193968.

30. Fernández-Calleja JMS, Bouwman LMS, Swarts HJM, Oosting A, Keijer J, van Schothorst EM. Extended indirect calorimetry with isotopic CO_2_ sensors for prolonged and continuous quantification of exogenous vs. total substrate oxidation in mice. Sci Rep. 2019 Aug 8;9(1):11507. doi: 10.1038/s41598-019-47977-w. PMID: 31395916; PMCID: PMC6687832.

31. Fischer AW, Csikasz RI, von Essen G, Cannon B, Nedergaard J. No insulating effect of obesity. Am J Physiol Endocrinol Metab. 2016 Jul 1;311(1):E202–13. doi: 10.1152/ajpendo.00093.2016. Epub 2016 May 17. PMID: 27189935.

32. Fiuza-Luces C, Santos-Lozano A, García-Silva MT, Martín-Hernández E, Quijada-Fraile P, Marín-Peiró M, Campos P, Arenas J, Lucía A, Martín MA, Morán M. Assessment of resting energy expenditure in pediatric mitochondrial diseases with indirect calorimetry. Clin Nutr. 2016 Dec;35(6):1484–1489. doi: 10.1016/j.clnu.2016.03.024. Epub 2016 Apr 7. PMID: 27105558.

33. Ganeshan K, Chawla A. Warming the mouse to model human diseases. Nat Rev Endocrinol. 2017 Aug;13(8):458–465. doi: 10.1038/nrendo.2017.48. Epub 2017 May 12. PMID: 28497813; PMCID: PMC5777302.

34. Gordon CJ. The mouse thermoregulatory system: Its impact on translating biomedical data to humans. Physiol Behav. 2017 Oct 1;179:55–66. doi: 10.1016/j.physbeh.2017.05.026. Epub 2017 May 19. PMID: 28533176; PMCID: PMC6196327.

35. Guzmán-Ruiz MA, Ramirez-Corona A, Guerrero-Vargas NN, Sabath E, Ramirez-Plascencia OD, Fuentes-Romero R, León-Mercado LA, Basualdo Sigales M, Escobar C, Buijs RM. Role of the Suprachiasmatic and Arcuate Nuclei in Diurnal Temperature Regulation in the Rat. J Neurosci. 2015 Nov 18;35(46):15419–29. doi: 10.1523/JNEUROSCI.1449-15.2015. PMID: 26586828; PMCID: PMC6605483.

36. Hanaford A, Johnson SC. The immune system as a driver of mitochondrial disease pathogenesis: a review of evidence. Orphanet J Rare Dis. 2022 Sep 2;17(1):335. doi: 10.1186/s13023-022-02495-3. PMID: 36056365; PMCID: PMC9438277.

37. Henke MT, Prigione A, Schuelke M. Disease models of Leigh syndrome: From yeast to organoids. J Inherit Metab Dis. 2024 Oct 9. doi: 10.1002/jimd.12804. Epub ahead of print. PMID: 39385390.

38. Henkel S, Frings-Meuthen P, Diekmann C, Coenen M, Stoffel-Wagner B, Németh R, Pesta D, Egert S. Influence of Ambient Temperature on Resting Energy Expenditure in Metabolically Healthy Males and Females. J Nutr. 2025 Mar;155(3):862–870. doi: 10.1016/j.tjnut.2025.01.013. Epub 2025 Jan 13. PMID: 39814140; PMCID: PMC11934286.

39. Hou Y, Xie Z, Zhao X, Yuan Y, Dou P, Wang Z. Appendicular skeletal muscle mass: A more sensitive biomarker of disease severity than BMI in adults with mitochondrial diseases. PLoS One. 2019 Jul 25;14(7):e0219628. doi: 10.1371/journal.pone.0219628. PMID: 31344055; PMCID: PMC6657836.

40. Hylander BL, Repasky EA. Thermoneutrality, Mice, and Cancer: A Heated Opinion. Trends Cancer. 2016 Apr;2(4):166–175. doi: 10.1016/j.trecan.2016.03.005. Epub 2016 Apr 22. PMID: 28741570.

41. Hylander BL, Qiao G, Cortes Gomez E, Singh P, Repasky EA. Housing temperature plays a critical role in determining gut microbiome composition in research mice: Implications for experimental reproducibility. Biochimie. 2023 Jul;210:71–81. doi: 10.1016/j.biochi.2023.01.016. Epub 2023 Jan 21. PMID: 36693616; PMCID: PMC10953156.

42. Jackson TC, Manole MD, Kotermanski SE, Jackson EK, Clark RS, Kochanek PM. Cold stress protein RBM3 responds to temperature change in an ultra-sensitive manner in young neurons. Neuroscience. 2015 Oct 1;305:268–78. doi: 10.1016/j.neuroscience.2015.08.012. Epub 2015 Aug 8. PMID: 26265550; PMCID: PMC4570027.

43. Jain IH, Zazzeron L, Goli R, Alexa K, Schatzman-Bone S, Dhillon H, Goldberger O, Peng J, Shalem O, Sanjana NE, Zhang F, Goessling W, Zapol WM, Mootha VK. Hypoxia as a therapy for mitochondrial disease. Science. 2016 Apr 1;352(6281):54–61. doi: 10.1126/science.aad9642. Epub 2016 Feb 25. PMID: 26917594; PMCID: PMC4860742.

44. James CM, Olejniczak SH, Repasky EA. How murine models of human disease and immunity are influenced by housing temperature and mild thermal stress. Temperature (Austin). 2022 Jul 15;10(2):166–178. doi: 10.1080/23328940.2022.2093561. PMID: 37332306; PMCID: PMC10274546.

45. John LM, Petersen N, Gerstenberg MK, Torz L, Pedersen K, Christoffersen BØ, Kuhre RE. Housing-temperature reveals energy intake counter-balances energy expenditure in normal-weight, but not diet-induced obese, male mice. Commun Biol. 2022 Sep 10;5(1):946. doi: 10.1038/s42003-022-03895-8. PMID: 36088386; PMCID: PMC9464191.

46. Johnson SC, Yanos ME, Kayser EB, Quintana A, Sangesland M, Castanza A, Uhde L, Hui J, Wall VZ, Gagnidze A, Oh K, Wasko BM, Ramos FJ, Palmiter RD, Rabinovitch PS, Morgan PG, Sedensky MM, Kaeberlein M. mTOR inhibition alleviates mitochondrial disease in a mouse model of Leigh syndrome. Science. 2013 Dec 20;342(6165):1524–8. doi: 10.1126/science.1244360. Epub 2013 Nov 14. PMID: 24231806; PMCID: PMC4055856.

47. Joko T, Iwashige K, Hashimoto T, Ono Y, Kobayashi K, Sekiguchi N, Kuroki T, Yanase T, Takayanagi R, Umeda F, Nawata H. A case of mitochondrial encephalomyopathy, lactic acidosis and stroke-like episodes associated with diabetes mellitus and hypothalamo-pituitary dysfunction. Endocr J. 1997 Dec;44(6):805–9. doi: 10.1507/endocrj.44.805. PMID: 9622295.

48. Jørgensen LB, Hansen AM, Willot Q, Overgaard J. Balanced mitochondrial function at low temperature is linked to cold adaptation in Drosophila species. J Exp Biol. 2023 Apr 15;226(8):jeb245439. doi: 10.1242/jeb.245439. Epub 2023 Apr 14. PMID: 36939380.

49. Kahlhöfer F, Kmita K, Wittig I, Zwicker K, Zickermann V. Accessory subunit NUYM (NDUFS4) is required for stability of the electron input module and activity of mitochondrial complex I. Biochim Biophys Acta Bioenerg. 2017 Feb;1858(2):175–181. doi: 10.1016/j.bbabio.2016.11.010. Epub 2016 Nov 19. PMID: 27871794.

50. Keijer J, Li M, Speakman JR. What is the best housing temperature to translate mouse experiments to humans? Mol Metab. 2019a Jul;25:168–176. doi: 10.1016/j.molmet.2019.04.001. Epub 2019 Apr 6. PMID: 31003945; PMCID: PMC6599456.

51. Keijer J, Li M, Speakman JR. To best mimic human thermal conditions, mice should be housed slightly below thermoneutrality. Mol Metab. 2019b Aug;26:4. doi: 10.1016/j.molmet.2019.05.007. Epub 2019 May 28. PMID: 31182417; PMCID: PMC6667391.

52. Kelly AM, Fricker BA, Wallace KJ. Protocol for multiplex fluorescent immunohistochemistry in free-floating rodent brain tissues. STAR Protoc. 2022 Dec 16;3(4):101672. doi: 10.1016/j.xpro.2022.101672. Epub 2022 Sep 14. PMID: 36107743; PMCID: PMC9483644.

53. Kempuraj D, Dourvetakis KD, Cohen J, Valladares DS, Joshi RS, Kothuru SP, Anderson T, Chinnappan B, Cheema AK, Klimas NG, Theoharides TC. Neurovascular unit, neuroinflammation and neurodegeneration markers in brain disorders. Front Cell Neurosci. 2024 Oct 25;18:1491952. doi: 10.3389/fncel.2024.1491952. PMID: 39526043; PMCID: PMC11544127.

54. Khumalo SG, Lindeque JZ, Venter M. Tissue-Specific Regulation of Fatty Acid Metabolism in a Mouse Model of Isolated Complex I Deficiency. Proteomics. 2025 Jun 1:e13969. doi: 10.1002/pmic.13969. Epub ahead of print. PMID: 40451765.

55. Koene S, Dirks I, van Mierlo E, de Vries PR, Janssen AJWM, Smeitink JAM, Bergsma A, Essers H, Meijer K, de Groot IJM. Domains of Daily Physical Activity in Children with Mitochondrial Disease: A 3D Accelerometry Approach. JIMD Rep. 2017;36:7–17. doi: 10.1007/8904_2016_35. Epub 2017 Jan 17. PMID: 28092092; PMCID: PMC5680282.

56. Koopman WJH, Distelmaier F, Smeitink JA, Willems PH. OXPHOS mutations and neurodegeneration. EMBO J. 2013 Jan 9;32(1):9–29. doi: 10.1038/emboj.2012.300. Epub 2012 Nov 13. PMID: 23149385; PMCID: PMC3545297.

57. Kozicz T, Morava E. Mitochondria as the key to understanding neuroinflammation. Brain. 2025 May 6:awaf173. doi: 10.1093/brain/awaf173. Epub ahead of print. PMID: 40326287.

58. Kozyreva, T.V. Adaptation to cold of homeothermic organism: changes in afferent and efferent links of the thermoregulatory system. J. Exp. Integr. Med. 2013; 3(4):255–265. DOI:10.5455/jeim.010813.ir.013

59. Kruse SE, Watt WC, Marcinek DJ, Kapur RP, Schenkman KA, Palmiter RD. Mice with mitochondrial complex I deficiency develop a fatal encephalomyopathy. Cell Metab. 2008 Apr;7(4):312–20. doi: 10.1016/j.cmet.2008.02.004. PMID: 18396137; PMCID: PMC2593686.

60. Lac M, Tavernier G, Moro C. Does housing temperature influence glucose regulation and muscle-fat crosstalk in mice? Biochimie. 2023 Jul;210:35–39. doi: 10.1016/j.biochi.2023.01.019. Epub 2023 Feb 8. PMID: 36758717.

61. Lamont RE, Beaulieu CL, Bernier FP, Sparkes R, Innes AM, Jackel-Cram C, Ober C, Parboosingh JS, Lemire EG. A novel NDUFS4 frameshift mutation causes Leigh disease in the Hutterite population. Am J Med Genet A. 2017 Mar;173(3):596–600. doi: 10.1002/ajmg.a.37983. Epub 2016 Sep 27. PMID: 27671926.

62. Lee C, Kim Y, Kaang BK. The Primary Motor Cortex: The Hub of Motor Learning in Rodents. Neuroscience. 2022 Mar 1;485:163–170. doi: 10.1016/j.neuroscience.2022.01.009. Epub 2022 Jan 17. PMID: 35051529.

63. Leigh D. Subacute necrotizing encephalomyelopathy in an infant. J Neurol Neurosurg Psychiatry. 1951 Aug;14(3):216–21. doi: 10.1136/jnnp.14.3.216. PMID: 14874135; PMCID: PMC499520.

64. Li D, Liang C, Zhang T, Marley JL, Zou W, Lian M, Ji D. Pathogenic mitochondrial DNA 3243A>G mutation: From genetics to phenotype. Front Genet. 2022 Oct 6;13:951185. doi: 10.3389/fgene.2022.951185. PMID: 36276941; PMCID: PMC9582660.

65. Martin-Perez M, Grillo AS, Ito TK, Valente AS, Han J, Entwisle SW, Huang HZ, Kim D, Yajima M, Kaeberlein M, Villén J. PKC downregulation upon rapamycin treatment attenuates mitochondrial disease. Nat Metab. 2020 Dec;2(12):1472–1481. doi: 10.1038/s42255-020-00319-x. Epub 2020 Dec 14. PMID: 33324011; PMCID: PMC8017771.

66. McCormick EM, Keller K, Taylor JP, Coffey AJ, Shen L, Krotoski D, Harding B; NICHD ClinGen U24 Mitochondrial Disease Gene Curation Expert Panel; Gai X, Falk MJ, Zolkipli-Cunningham Z, Rahman S. Expert Panel Curation of 113 Primary Mitochondrial Disease Genes for the Leigh Syndrome Spectrum. Ann Neurol. 2023 Oct;94(4):696–712. doi: 10.1002/ana.26716. Epub 2023 Aug 12. PMID: 37255483; PMCID: PMC10763625.

67. MacDonald CR, Choi JE, Hong CC, Repasky EA. Consideration of the importance of measuring thermal discomfort in biomedical research. Trends Mol Med. 2023 Aug;29(8):589–598. doi: 10.1016/j.molmed.2023.05.010. Epub 2023 Jun 15. PMID: 37330365; PMCID: PMC10619709.

68. Mei J, Riedel N, Grittner U, Endres M, Banneke S, Emmrich JV. Body temperature measurement in mice during acute illness: implantable temperature transponder versus surface infrared thermometry. Sci Rep. 2018 Feb 23;8(1):3526. doi: 10.1038/s41598-018-22020-6. PMID: 29476115; PMCID: PMC5824949.

69. Müller TD, Klingenspor M, Tschöp MH. Revisiting energy expenditure: how to correct mouse metabolic rate for body mass. Nat Metab. 2021 Sep;3(9):1134–1136. doi: 10.1038/s42255-021-00451-2.

70. Muñoz Zamora A, Douglas A, Conway PB, Urrieta E, Moniz T, O’Leary JD, Marks L, Denny CA, Ortega-de San Luis C, Lynch L, Ryan TJ. Cold memories control whole-body thermoregulatory responses. Nature. 2025 Apr 23. doi: 10.1038/s41586-025-08902-6. Epub ahead of print. PMID: 40269165.

71. Naveed M, Smedlund K, Zhou QG, Cai W, Hill JW. Astrocyte involvement in metabolic regulation and disease. Trends Endocrinol Metab. 2025 Mar;36(3):219–234. doi: 10.1016/j.tem.2024.08.001. Epub 2024 Aug 29. PMID: 39214743; PMCID: PMC11868460.

72. Nakamura K. Central circuitries for body temperature regulation and fever. Am J Physiol Regul Integr Comp Physiol. 2011 Nov;301(5):R1207–28. doi: 10.1152/ajpregu.00109.2011. Epub 2011 Sep 7. PMID: 21900642.

73. Noorda G, van Achterberg T, van der Hooft T, Smeitink JAM, Schoonhoven L, van Engelen B. Problems of adults with a mitochondrial disease - the patients’ perspective: focus on loss. JIMD Rep. 2012;6:85–94. doi: 10.1007/8904_2011_121. Epub 2012 Feb 24. PMID: 23430944; PMCID: PMC3565683.

74. Ortigoza-Escobar JD, Oyarzabal A, Montero R, Artuch R, Jou C, Jiménez C, Gort L, Briones P, Muchart J, López-Gallardo E, Emperador S, Pesini ER, Montoya J, Pérez B, Rodríguez-Pombo P, Pérez-Dueñas B. Ndufs4 related Leigh syndrome: A case report and review of the literature. Mitochondrion. 2016 May;28:73–8. doi: 10.1016/j.mito.2016.04.001. Epub 2016 Apr 11. PMID: 27079373.

75. Pamenter ME, Lau GY, Richards JG. Effects of cold on murine brain mitochondrial function. PLoS One. 2018 Dec 6;13(12):e0208453. doi: 10.1371/journal.pone.0208453. PMID: 30521596; PMCID: PMC6283463.

76. Papa S, Sardanelli AM, Cocco T, Speranza F, Scacco SC, Technikova-Dobrova Z. The nuclear-encoded 18 kDa (IP) AQDQ subunit of bovine heart complex I is phosphorylated by the mitochondrial cAMP-dependent protein kinase. FEBS Lett. 1996 Feb 5;379(3):299–301. doi: 10.1016/0014-5793(95)01532-9. PMID: 8603710.

77. Quintana A, Kruse SE, Kapur RP, Sanz E, Palmiter RD. Complex I deficiency due to loss of Ndufs4 in the brain results in progressive encephalopathy resembling Leigh syndrome. Proc Natl Acad Sci U S A. 2010 Jun 15;107(24):10996–1001. doi: 10.1073/pnas.1006214107. Epub 2010 Jun 1. PMID: 20534480; PMCID: PMC2890717.

78. Quintana A, Zanella S, Koch H, Kruse SE, Lee D, Ramirez JM, Palmiter RD. Fatal breathing dysfunction in a mouse model of Leigh syndrome. J Clin Invest. 2012 Jul;122(7):2359–68. doi: 10.1172/JCI62923. Epub 2012 Jun 1. PMID: 22653057; PMCID: PMC3387817.

79. Rahman S. Leigh syndrome. Handb Clin Neurol. 2023;194:43–63. doi: 10.1016/B978-0-12-821751-1.00015-4. PMID: 36813320.

80. Rahman S. Complex I deficiency remains the most frequent cause of Leigh syndrome spectrum. Brain Commun. 2024 Dec 23;7(1):fcae470. doi: 10.1093/braincomms/fcae470. PMID: 39816196; PMCID: PMC11733768.

81. Rahman S, Thorburn D. Nuclear Gene-Encoded Leigh Syndrome Spectrum Overview. 2015 Oct 1 [updated 2020 Jul 16]. In: Adam MP, Feldman J, Mirzaa GM, Pagon RA, Wallace SE, Amemiya A, editors. GeneReviews [Internet]. Seattle (WA): University of Washington, Seattle; 1993–2024. PMID: 26425749.

82. Rango M, Arighi A, Bonifati C, Del Bo R, Comi G, Bresolin N. The brain is hypothermic in patients with mitochondrial diseases. J Cereb Blood Flow Metab. 2014 May;34(5):915–20. doi: 10.1038/jcbfm.2014.38. Epub 2014 Mar 12. PMID: 24619278; PMCID: PMC4013774.

83. Rosenthal LM, Leithner C, Tong G, Streitberger KJ, Krech J, Storm C, Schmitt KRL. RBM3 and CIRP expressions in targeted temperature management treated cardiac arrest patients-A prospective single center study. PLoS One. 2019 Dec 10;14(12):e0226005. doi: 10.1371/journal.pone.0226005. PMID: 31821351; PMCID: PMC6903712.

84. Sadler DG, Treas L, Sikes JD, Porter C. A modest change in housing temperature alters whole body energy expenditure and adipocyte thermogenic capacity in mice. Am J Physiol Endocrinol Metab. 2022 Dec 1;323(6):E517–E528. doi: 10.1152/ajpendo.00079.2022. Epub 2022 Nov 9. PMID: 36351253; PMCID: PMC9744648.

85. Schirris TJJ, Rossell S, de Haas R, Frambach SJCM, Hoogstraten CA, Renkema GH, Beyrath JD, Willems PHGM, Huynen MA, Smeitink JAM, Russel FGM, Notebaart RA. Stimulation of cholesterol biosynthesis in mitochondrial complex I-deficiency lowers reductive stress and improves motor function and survival in mice. Biochim Biophys Acta Mol Basis Dis. 2021 Apr 1;1867(4):166062. doi: 10.1016/j.bbadis.2020.166062. Epub 2021 Jan 13. PMID: 33385517.

86. Seeley RJ, MacDougald OA. Mice as experimental models for human physiology: when several degrees in housing temperature matter. Nat Metab. 2021 Apr;3(4):443–445. doi: 10.1038/s42255-021-00372-0. PMID: 33767444; PMCID: PMC8987294.

87. Shil SK, Kagawa Y, Umaru BA, Nanto-Hara F, Miyazaki H, Yamamoto Y, Kobayashi S, Suzuki C, Abe T, Owada Y. Ndufs4 ablation decreases synaptophysin expression in hippocampus. Sci Rep. 2021 May 26;11(1):10969. doi: 10.1038/s41598-021-90127-4. PMID: 34040028; PMCID: PMC8155116.

88. Shin YC, Latorre-Muro P, Djurabekova A, Zdorevskyi O, Bennett CF, Burger N, Song K, Xu C, Paulo JA, Gygi SP, Sharma V, Liao M, Puigserver P. Structural basis of respiratory complex adaptation to cold temperatures. Cell. 2024 Nov 14;187(23):6584-6598.e17. doi: 10.1016/j.cell.2024.09.029. Epub 2024 Oct 11. PMID: 39395414; PMCID: PMC11601890.

89. Škop V, Liu N, Xiao C, Stinson E, Chen KY, Hall KD, Piaggi P, Gavrilova O, Reitman ML. Beyond day and night: The importance of ultradian rhythms in mouse physiology. Mol Metab. 2024 Jun;84:101946. doi: 10.1016/j.molmet.2024.101946. Epub 2024 Apr 23. PMID: 38657735; PMCID: PMC11070603.

90. Smeitink JAM, van den Heuvel L, DiMauro S. The genetics and pathology of oxidative phosphorylation. Nat Rev Genet. 2001 May;2(5):342–52. doi: 10.1038/35072063. PMID: 11331900.

91. Sofou K, De Coo IF, Isohanni P, Ostergaard E, Naess K, De Meirleir L, Tzoulis C, Uusimaa J, De Angst IB, Lönnqvist T, Pihko H, Mankinen K, Bindoff LA, Tulinius M, Darin N. A multicenter study on Leigh syndrome: disease course and predictors of survival. Orphanet J Rare Dis. 2014 Apr 15;9:52. doi: 10.1186/1750-1172-9-52. PMID: 24731534; PMCID: PMC4021638.

92. Speakman JR, Keijer J. Not so hot: Optimal housing temperatures for mice to mimic the thermal environment of humans. Mol Metab. 2012 Nov 8;2(1):5–9. doi: 10.1016/j.molmet.2012.10.002. PMID: 24024125; PMCID: PMC3757658.

93. Stenton SL, Zou Y, Cheng H, Liu Z, Wang J, Shen D, Jin H, Ding C, Tang X, Sun S, Han H, Ma Y, Zhang W, Jin R, Wang H, Sun D, Lv JL, Prokisch H, Fang F. Leigh Syndrome: A Study of 209 Patients at the Beijing Children’s Hospital. Ann Neurol. 2022 Apr;91(4):466–482. doi: 10.1002/ana.26313. Epub 2022 Mar 6. PMID: 35094435.

94. Stokes JC, Bornstein RL, James K, Park KY, Spencer KA, Vo K, Snell JC, Johnson BM, Morgan PG, Sedensky MM, Baertsch NA, Johnson SC. Leukocytes mediate disease pathogenesis in the Ndufs4(KO) mouse model of Leigh syndrome. JCI Insight. 2022 Mar 8;7(5):e156522. doi: 10.1172/jci.insight.156522. PMID: 35050903; PMCID: PMC8983133.

95. Tan CL, Knight ZA. Regulation of Body Temperature by the Nervous System. Neuron. 2018 Apr 4;98(1):31–48. doi: 10.1016/j.neuron.2018.02.022. PMID: 29621489; PMCID: PMC6034117.

96. Tansey EA, Johnson CD. Recent advances in thermoregulation. Adv Physiol Educ. 2015 Sep;39(3):139–48. doi: 10.1152/advan.00126.2014. PMID: 26330029.

97. Terburgh K, Coetzer J, Lindeque JZ, van der Westhuizen FH, Louw R. Aberrant BCAA and glutamate metabolism linked to regional neurodegeneration in a mouse model of Leigh syndrome. Biochim Biophys Acta Mol Basis Dis. 2021 May 1;1867(5):166082. doi: 10.1016/j.bbadis.2021.166082. Epub 2021 Jan 22. PMID: 33486097.

98. Thorburn DR, Rahman J, Rahman S. Mitochondrial DNA-Associated Leigh Syndrome and NARP. 2003 Oct 30 [updated 2023 May 4]. In: Adam MP, Feldman J, Mirzaa GM, Pagon RA, Wallace SE, Bean LJH, Gripp KW, Amemiya A, editors. GeneReviews [Internet]. Seattle (WA): University of Washington, Seattle; 1993–2024. PMID: 20301352.

99. Tong G, Endersfelder S, Rosenthal LM, Wollersheim S, Sauer IM, Bührer C, Berger F, Schmitt KR. Effects of moderate and deep hypothermia on RNA-binding proteins RBM3 and CIRP expressions in murine hippocampal brain slices. Brain Res. 2013 Apr 4;1504:74–84. doi: 10.1016/j.brainres.2013.01.041. Epub 2013 Feb 8. PMID: 23415676.

100. Tran LT, Park S, Kim SK, Lee JS, Kim KW, Kwon O. Hypothalamic control of energy expenditure and thermogenesis. Exp Mol Med. 2022 Apr;54(4):358–369. doi: 10.1038/s12276-022-00741-z. Epub 2022 Mar 17. PMID: 35301430; PMCID: PMC9076616.

101. Tschöp MH, Speakman JR, Arch JR, Auwerx J, Brüning JC, Chan L, Eckel RH, Farese RV Jr, Galgani JE, Hambly C, Herman MA, Horvath TL, Kahn BB, Kozma SC, Maratos-Flier E, Müller TD, Münzberg H, Pfluger PT, Plum L, Reitman ML, Rahmouni K, Shulman GI, Thomas G, Kahn CR, Ravussin E. A guide to analysis of mouse energy metabolism. Nat Methods. 2011 Dec 28;9(1):57–63. doi: 10.1038/nmeth.1806. PMID: 22205519; PMCID: PMC3654855.

102. van de Wal MAE, Adjobo-Hermans MJW, Keijer J, Schirris TJJ, Homberg JR, Wieckowski MR, Grefte S, van Schothorst EM, van Karnebeek C, Quintana A, Koopman WJH. Ndufs4 knockout mouse models of Leigh syndrome: pathophysiology and intervention. Brain. 2022 Mar 29;145(1):45–63. doi: 10.1093/brain/awab426. PMID: 34849584; PMCID: PMC8967107.

103. van de Wal MAE, Doornbos C, Bibbe JM, Homberg JR, van Karnebeek C, Huynen MA, Keijer J, van Schothorst EM, ‘t Hoen PAC, Janssen MCH, Adjobo-Hermans MJW, Wieckowski MR, Koopman WJH. Ndufs4 knockout mice with isolated complex I deficiency engage a futile adaptive brain response. Biochim Biophys Acta Proteins Proteom. 2025 Oct 11;1873(1):141055. doi: 10.1016/j.bbapap.2024.141055. Epub ahead of print. PMID: 39395749.

104. van den Heuvel L, Ruitenbeek W, Smeets R, Gelman-Kohan Z, Elpeleg O, Loeffen J, Trijbels F, Mariman E, de Bruijn D, Smeitink JAM. Demonstration of a new pathogenic mutation in human complex I deficiency: a 5-bp duplication in the nuclear gene encoding the 18-kD (AQDQ) subunit. Am J Hum Genet. 1998 Feb;62(2):262–8. doi: 10.1086/301716. PMID: 9463323; PMCID: PMC1376892.

105. Von Schulze AT, Deng F, Fuller KNZ, Franczak E, Miller J, Allen J, McCoin CS, Shankar K, Ding WX, Thyfault JP, Geiger PC. Heat Treatment Improves Hepatic Mitochondrial Respiratory Efficiency via Mitochondrial Remodeling. Function (Oxf). 2021 Jan 22;2(2):zqab001. doi: 10.1093/function/zqab001. PMID: 33629069; PMCID: PMC7886620.

106. Yagasaki H, Ohyama T, Narusawa H, Nakane T. Hypothermic reaction after infection in an infant with pyruvate dehydrogenase complex deficiency. Pediatr Neonatol. 2019 Aug;60(4):475–476. doi: 10.1016/j.pedneo.2019.04.005. Epub 2019 Apr 13. PMID: 31064704.

107. Yin Z, Agip AA, Bridges HR, Hirst J. Structural insights into respiratory complex I deficiency and assembly from the mitochondrial disease-related ndufs4^-/-^mouse. EMBO J. 2024 Jan;43(2):225–249. doi: 10.1038/s44318-023-00001-4. Epub 2024 Jan 2. PMID: 38177503; PMCID: PMC10897435.

108. Zhao Z, Yang R, Li M, Bao M, Huo D, Cao J, Speakman JR. Effects of ambient temperatures between 5 and 35 °C on energy balance, body mass and body composition in mice. Mol Metab. 2022a Oct;64:101551. doi: 10.1016/j.molmet.2022.101551. Epub 2022 Jul 20. PMID: 35870706; PMCID: PMC9382332.

109. Zhao Z, Cao J, Niu C, Bao M, Xu J, Huo D, Liao S, Liu W, Speakman JR. Body temperature is a more important modulator of lifespan than metabolic rate in two small mammals. Nat Metab. 2022b Mar;4(3):320–326. doi: 10.1038/s42255-022-00545-5. Epub 2022 Mar 14. PMID: 35288719.

110. Zweers HEE, Janssen MCH, Wanten GJA. Optimal Estimate for Energy Requirements in Adult Patients With the m.3243A>G Mutation in Mitochondrial DNA. JPEN J Parenter Enteral Nutr. 2021 Jan;45(1):158–164. doi: 10.1002/jpen.1965. Epub 2020 Aug 1. PMID: 32696575; PMCID: PMC7891583.

